# Leveraging weighted quartet distributions for enhanced species tree inference from genome-wide data

**DOI:** 10.1101/2024.09.15.613103

**Authors:** Navid Bin Hasan, Avijit Biswas, Zahin Wahab, Mahim Mahbub, Rezwana Reaz, Md Shamsuzzoha Bayzid

## Abstract

Species tree estimation from genes sampled from throughout the whole genome is challenging in the presence of gene tree discordance, often caused by incomplete lineage sorting (ILS), where alleles can coexist in populations for periods that may span several speciation events. Quartet-based summary methods for estimating species trees from a collection of gene trees are becoming popular due to their high accuracy and theoretical guarantees of robustness to arbitrarily high amounts of ILS. ASTRAL, the most widely used quartet-based method, aims to infer species trees by maximizing the number of quartets in the gene trees that are consistent with the species tree. An alternative approach (as in wQFM) is to infer quartets for all subsets of four species and amalgamate them into a coherent species tree. While summary methods can be highly sensitive to gene tree estimation errors–especially when gene trees are derived from short alignments–quartet amalgamation offers an advantage by potentially bypassing the need for gene tree estimation. However, greatly understudied is the choice of weighted quartet inference method and downstream effects on species tree estimations under realistic model conditions. In this study, we investigated a broad range of methods for generating weighted quartets and critically assessed their impact on species tree inference. Our results on a collection of simulated and empirical datasets suggest that amalgamating quartets weighted based on gene tree frequencies (GTF) typically produces more accurate trees than leading quartet-based methods like ASTRAL and SVDquartets. Further enhancements in GTF-based weighted quartet estimation were achieved by accounting for gene tree uncertainty, through the utilization of a distribution of trees for each gene (instead of a single tree), by employing traditional nonparametric bootstrapping methods or Bayesian MCMC sampling. Our study provides evidence that the careful generation and amalgamation of weighted quartets, as implemented in methods like wQFM, can lead to significantly more accurate trees compared to widely employed methods like ASTRAL, especially in the face of gene tree estimation errors.

## 1 Introduction

The estimation of species trees using multiple loci has become increasingly common. However, species tree estimation from multi-locus data sampled from throughout the whole genome is difficult because different loci can have different phylogenetic histories, a phenomenon known as *gene tree discordance* which occurs due to several different biological processes, including Incomplete Lineage Sorting (ILS), Gene Duplication (GD), and Horizontal Gene Transfer (HGT) [1, 2]. In particular, many groups of species evolve with rapid speciation events, a process that tends to create conflicts between gene trees and species trees due to ILS [3], which is modeled by the multi-species coalescent model [4].

In the presence of gene tree discordance, standard methods for estimating species trees, such as concatenation (which concatenates multiple sequence alignments of different genes into a single supermatrix, which is then used to estimate the species tree), can be statistically inconsistent [5, 6], and produce incorrect trees with high support [7]. Therefore, a two-step process, where gene trees are first inferred independently from sequence data and then combined using “summary methods” to estimate species trees by summarizing the input gene trees, is becoming increasingly popular, and many of them are provably statistically consistent under ILS [8–17].

Quartet-based summary methods have gained significant attention because quartets (4-leaf unrooted gene trees) avoid the “anomaly zone” – a scenario where the most likely gene tree topology may differ from the species tree topology [18–20]. ASTRAL, the most widely used summary method, takes a set of gene trees to infer a species tree that maximizes the number of quartets in the gene trees that are consistent with the species tree. Another class of methods, including wQFM and wQMC (known as quartet amalgamation techniques), involves inferring individual quartets for each set of four taxa and then combining them into a cohesive species tree. A major challenge in summarizing gene trees is that when we infer gene trees, often from relatively short sequences, the estimated gene trees tend to be highly error-prone – making summary methods sensitive to gene tree estimation error. A broader impact and a clear benefit of the quartet amalgamation based approach over ASTRAL is that it can be used outside the context of gene tree estimation. Chifman and Kubatko introduced SVDquartets [13], a popular quartet-based method, which avoids estimating trees on each locus and hence bypasses the issue of gene tree estimation error. It combines multi-locus unlinked single-site data, infers the quartet trees for all subsets of four species, and then combines the set of quartet trees into a species tree using a quartet amalgamation heuristic such as QFM [10] or QMC [21]. There is ample evidence that assigning weights to quartets – where the weight of a quartet denotes the relative confidence of a particular quartet topology out of the three alternate topologies on a set of four taxa – can enhance phylogenetic analyses [9, 12] despite the presence of gene tree estimation error. Zhang et al. showed that wASTRAL, the variant of ASTRAL that assigns weights to quartets based on gene tree uncertainty, outperforms the unweighted version of ASTRAL on simulated data in terms of its topology and branch support values [22]. Casanellas et al. proposed a new weighting system for quartets based on algebraic and semi-algebraic tools [23], which is specially tailored to deal with data evolving under heterogeneous rates.

Despite the popularity of quartet-based methods and the growing awareness that appropriate weighting schemes of quartets can improve phylogenomic analyses, a greatly understudied aspect is the way weighted quartets are estimated before summarizing/amalgamating them. In this study, we address this gap by computing weighted quartets using a broad range of meaningful ways, using widely used Bayesian, maximum likelihood (ML), and statistical tools, including MrBayes [24], BUCKy [25,26], RAxML [27], and SVDquartets [13]. We compare these weighted quartet generation strategies that significantly differ in their underlying statistical frameworks and computational demands. We combine these various quartet distribution generation methods with popular quartet amalgamation techniques like wQFM and wQMC. We performed an extensive experimental study and compared these techniques with the leading species tree estimation methods such as ASTRAL, BUCKy, and SVDquartets. Specifically, our study addresses the following research questions (RQs).

1. RQ1: Propose and explore a wide range of ways to generate quartet distributions and assess their performance in terms of species tree accuracy.
2. RQ2: Investigate the relative performance of the most popular quartet amalgamation techniques, wQFM and wQMC, when paired with various quartet distribution generation approaches.
3. RQ3: Compare the relative performance of the promising approaches identified in RQ1 and RQ2 with leading quartet-based species tree estimation methods (e.g., ASTRAL, SVDquar-tets, and BUCKy).
4. RQ4: Determine if the quartet scores of species trees (defined to be the number of quartets from the set of input gene trees that agree with the species tree) estimated by statistically consistent methods are predictive of species tree accuracy under practical model conditions (i.e., limited number of gene trees and low phylogenetic signal per gene).
5. RQ5: Evaluate the performance of the best methods identified in RQ1-RQ3 on real biological datasets.

## 2 Experiment design

### 2.1 Methods

In our study, we utilized leading quartet-based summary methods to estimate species trees using gene trees, multiple sequence alignments (MSAs), or a combination of both. Given that prior studies have already compared summary methods with combined analysis (CA) and established its relative performance [8, 28, 29], we chose not to include CA in our experiments. The methods included in our study are as follows:

#### ASTRAL and wASTRAL

ASTRAL is a statistically consistent method under the MSC model that tries to solve the Maximum Quartet Support Species Tree (MQSST) problem [8]. It takes a set of gene trees as input and seeks to find a species tree so that the number of induced quartets in the gene trees that are consistent with the species tree is maximized. We analyzed both unweighted ASTRAL and weighted ASTRAL (wASTRAL) [22].

#### wQFM and wQMC

These are two highly accurate methods for species tree estimation that amalgamate weighted quartets. wQFM and wQMC extend the popular (unweighted) quartet amalgamation techniques QFM and QMC to a weighted setting.

#### SVDquartets

The SVDquartets algorithm takes unlinked multi-locus data as input and, for a set of four taxa (*a, b, c, d*), assigns a score to each of the three possible quartet topologies (*ab*|*cd, ac*|*bd, ad*|*bc*) using algebraic statistics and singular value decomposition (SVD). The quartet topology, among the three alternative topologies with the lowest “SVD score”, is selected as the “dominant” or true topology for that quartet. These selected quartets are then amalgamated using quartet assembly techniques, such as QFM or QMC, to construct a species tree.

#### BUCKy

Given a distribution of trees for each gene (usually generated by MrBayes), BUCKy uses Bayesian concordance analysis to estimate the Concordance Factors (CF) of quartets. These CFs are used to estimate a concordance tree (BUCKY-con) and a population tree (BUCKY-pop). We use the population tree in our analyses, as this is provably statistically consistent. We estimated gene tree distributions using MrBayes (MB) and using RAxML with bootstrapping as input to BUCKy, denoted by BUCKy-MB and BUCKy-RAxML, respectively. For various datasets and model conditions analyzed in this study, we ran BUCKy using a sufficiently large number of MCMC iterations to reach sufficiently low standard deviations for the concordance factors to suggest possible convergence.

wQFM and wQMC, in general, can be used to amalgamate any set of given weighted quartets. Therefore, the broader impact and a clear benefit of wQFM and wQMC over ASTRAL and other similar summary methods is that these can be used outside the context of gene tree estimation. In this study, we have proposed and investigated a wide range of ways for computing weighted quartets and their impact on species tree inference when coupled with wQFM and wQMC. As a result, we have explored a large number of combinations with different quartet generation techniques and quartet amalgamation techniques. We define these methods in Table 1 for wQFM (using the convention wQFM–⟨quartet-generation-technique⟩), but all of them are applicable for wQMC. Figure 1 shows a schematic diagram of our experimental study.

**Figure 1:**
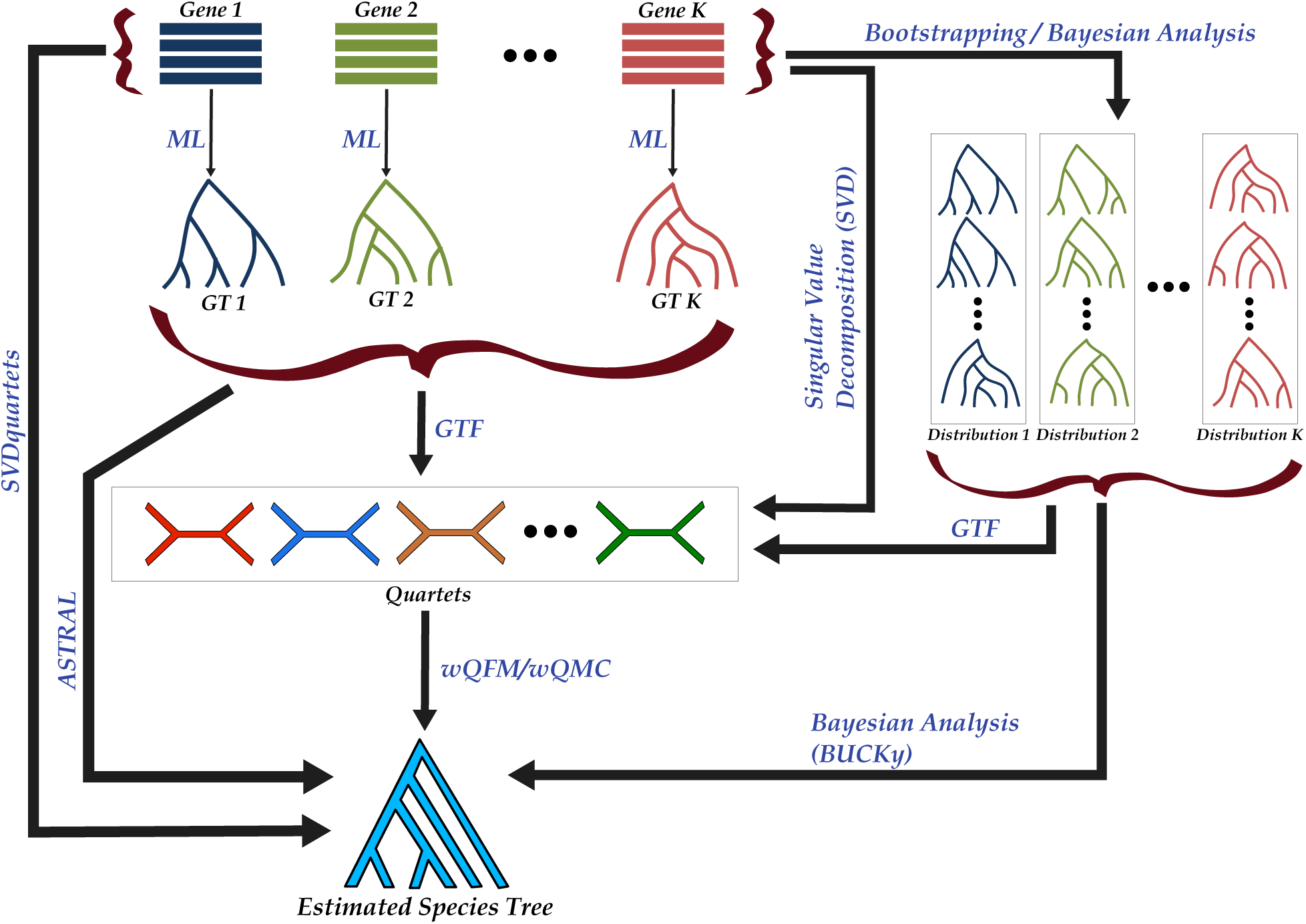
Schematic diagram of the experimental study pipeline. The process begins with a set of *k* input gene sequences, from which *k* maximum likelihood (ML) gene trees are estimated using RAxML. In addition to ML gene trees, distributions of trees for each gene is estimated using bootstrapping (RAxML) or Bayesian methods (MrBayes). Weighted quartet distributions are then generated from these sets of gene trees, with the quartet’s weight determined by its frequency across the input trees. Furthermore, SVDquartets is used to directly infer a weighted quartet distribution from the concatenated gene sequences. Finally, wQFM or wQMC amalgamates the various quartet sets to estimate species trees. For comparison, species trees are also inferred from gene trees using ASTRAL, from gene tree distributions using BUCKy, and from concatenated gene sequences using SVDquartets.

**Table 1:** Brief descriptions of the quartet generation strategies explored in this study. These quartets are amalgamated by wQFM and wQMC to estimate species trees.

| Method | Description |
| --- | --- |
| wQFM-GTF-all | wQFM-GTF refers to wQFM with quartets weighted based on gene tree frequencies (GTF), which is the number of occurrences of a quartet in the input gene trees. wQFM-GTF-all considers all three alternative topologies for each set of four taxa. Therefore, unlike SVDquartets and BUCKy-pop, which consider only the dominant quartet for each set of four taxa, wQFM-GTF-all computes weights for every possible four-leaf tree (and so for each of the three possible unrooted trees for every four leaves), and then combines this set of weighted quartet trees into a tree on the full set of species. |
| wQFM-GTF-dom | This method considers only the dominant quartets (i.e., the quartet topology with the highest gene tree frequency) and their weights for each set of four taxa. |
| wQFM-GTF-dom<br>(without weights) | Similar to wQFM-GTF-dom, only the dominant quartets are taken into consideration, but weights of the quartets are ignored (i.e., each dominant quartet is assigned unit weight). |
| wQFM-GTF-RAxML-BS | This approach refers to wQFM-GTF-all when run on bootstrap (BS) gene tree distributions for every gene computed by RAxML. We use nonparametric bootstrapping using RAxML to produce 200 trees for each gene. We then count GTF for all possible quartets based on all these BS trees. The motivation behind using a BS gene tree distribution instead of single best tree estimate for each gene is that a larger pool of trees is likely to augment the quartets with more meaningful weights and better address gene tree uncertainty. |
| wQFM-GTF-MB | Similar to wQFM-GTF-RAxML-BS, but here the set of trees for each gene is computed using MrBayes (by performing Bayesian analysis on its MSA) instead of RAxML. MrBayes was run twice, each for 55000 generations, and we sample a total of 200 gene trees, excluding burn-in samples. |
| wQFM-SVD-Exp | SVDquartets has two modes for generating weighted quartets: i) exponential and ii) reciprocal. wQFM-SVD-Exp refers to wQFM when run on weighted quartets computed by SVDquartets using the exponential weighting scheme. |
| wQFM-SVD-Rec | Similar to wQFM-SVD-Exp, but the weights are computed using the reciprocal weighting format. |
| wQFM-RAxML | It refers to wQFM with quartets weighted based on the likelihood scores computed by RAxML (using the fast quartet calculator option) given the combined alignment of the gene sequences. |
| wQFM-BUCKy-MB | wQFM with quartets weighted based on CFs, generated by BUCKy when a set of trees generated by MrBayes (MB) is provided as input. As the concordance factor of a quartet reflects the percentage of gene trees that support it, it is a natural choice as a weight for that quartet. |
| wQFM-BUCKy-RAxML | Similar to wQFM-BUCKy-MB except that the input gene trees to BUCKy are generated by RAxML bootstrapping instead of MrBayes. |

### 2.2 Dataset

#### Simulated dataset

We used previously studied simulated and real biological datasets to eval-uate the performance of various ways for generating weighted quartets (Table 1), and various quartet-based summary methods. We analyzed three small to moderate size dataset: the 11-, 15-and 37-taxon simulated datasets from [30, 31]. We did not include large datasets with hundreds of taxa in this study due to the computationally intensive nature of methods such as MrBayes and BUCKy, which require prohibitively long runtimes on larger datasets.

The 37-taxon mammalian simulated dataset was generated using the species tree estimated by MP-EST on the biological dataset studied in Song et al. [32]. This species tree, with branch lengths in coalescent units, served as the basis for producing a set of gene trees under the coalescent model. Consequently, the model tree exhibits an ILS level consistent with a coalescent analysis of the biological mammalian dataset, and other simulation properties reflect the biological sequences studied. We examined the impact of varying the number of genes (ranging from 25 to 500) and the degree of gene tree estimation error, which was controlled by adjusting the sequence length of the markers (from 250 bp to 1500 bp). Additionally, the levels of ILS were varied by modifying all internal branch lengths in the model species tree, either doubling or halving them. This resulted in three model conditions: 1X (moderate ILS), 0.5X (high ILS), and 2X (low ILS).

We also analyzed the high-ILS 11-taxon datasets from [30], but did not include the lower ILS model conditions as they are relatively easy to analyze [28]. These datasets vary in the number of genes and the extent of gene tree estimation error. Additionally, we examined 15-taxon datasets that also exhibit high levels of ILS and vary in both sequence lengths and the number of genes. Consequently, these simulated datasets offer a broad spectrum of challenging and practical model conditions, allowing us to rigorously evaluate the performance of various methods explored in this study.

#### Biological dataset

We reanalyzed the 37-taxon mammalian dataset from Song *et al.* [32] containing 447 genes across 37 mammals after removing 21 mislabeled genes (confirmed by the authors), and two other outlier genes. We also analyze two biological datasets: the mammalian dataset containing 447 genes across 37 taxa from Song *et al.* [32] and the avian dataset by Jarvis *et al.* [3] containing genome data of 48 avian species spanning most orders of birds. The avian dataset includes 14,446 loci comprising exons, introns, and UCEs.

### 2.3 Evaluation criteria

On the simulated datasets, we compared the estimated trees with the model species tree using normalized Robinson-Foulds (RF) distance [33]. The RF distance between two trees is defined as the sum of the bipartitions (splits) present in one tree but absent in the other, and vice versa. All the estimated trees in this study are binary, making False Positive (FP), False Negative (FN), and RF rates identical. Additionally, we compared the quartet scores (the number of quartets from the set of input gene trees that agree with the candidate species tree) of the trees estimated by different methods. For the biological dataset, we compared the estimated species trees with the established evolutionary relationships reported in the scientific literature. Multiple replicates of data for various model conditions were analyzed, and a two-sided Wilcoxon signed-rank test (with *α* = 0.05) was performed to measure the statistical significance of the differences between methods.

## 3 Results and discussion

In this section, we present the findings related to the research questions (RQs) explored in this study. For each research question, we compare the methods that we consider most appropriate for addressing that specific question.

### 3.1 RQ1: Finding the most appropriate quartet distributions

In RQ1, we have four separate experiments to assess the impact of different strategies for generating quartet distributions on species tree estimation.

- Experiment 1: What is the most effective approach for generating gene tree frequency (GTF)-based quartet distributions: using only dominant quartets (with or without weights) or incorporating all quartets??
- Experiment 2: Should we rely solely on the best maximum likelihood (bestML) gene trees for generating weighted quartets, or should we incorporate distributions of gene trees, using non-parametric bootsrapping or Bayesian MCMC sampling?
- Experiment 3: Does the weighted setting of SVDquartets lead to improved phylogenies?
- Experiment 4: How do ML-based and BUCKy-based quartet distributions compare to GTF-based quartet distributions??

#### Experiment 1: Dominant quartets vs all quartets

The basic idea of Combining Dominant Quartet Trees (CDQT) is to take the input set of gene trees G, compute a dominant quartet tree (i.e., the most frequent quartet topology) for every four species, and then combine the dominant quartet trees into a supertree on the full set of species using a preferred quartet amalgamation technique (QFM or QMC). CDQT is a statistically consistent approach [9]. We consider the dominant quartets with and without weights, where the weight of a quartet *q* is computed based on the number of *gt* ∈ GT that induce quartet topology *q*. These strategies are denoted by GTF-dom and GTF-dom (without weights), respectively. Another statistically consistent approach is *Weighted Maximum Quartet Consistency* (WMQC) problem [9], which computes weights for every possible quartet trees, and then combines this set of weighted quartet trees into a tree on the full set of species. Thus, it uses all possible quartets (i.e., 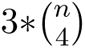 quartets for n taxa) with relative weights and is denoted by GTF-all. We analyzed these three different quartet distribution generation strategies (i.e., GTF-dom, GTF-dom (without weights), and GTF-all) using two leading weighted quartet amalgamation techniques, wQFM and wQMC.

The performance of six methods (three quartet generation strategies coupled with both wQFM and wQMC) on the 15-taxon dataset is shown in Figure 2, where we investigated the performance on varying gene tree estimation errors using 100bp and 1000bp sequence lengths and on varying numbers of genes (100 and 1000). Across all model conditions, except for the easiest one (1000bp-1000gt), where all methods perform very well, GTF-all is significantly better than GTF-dom – suggesting that using all quartets is better than using only the dominant ones. Moreover, as shown in previous studies [9], wQFM is, in general, more accurate than wQMC. Interestingly, wQMC achieved similar accuracies with dominant quartets both with and without weights. In contrast, wQFM performed better with weighted dominant quartets when the number of genes was relatively small but performed better without weights as the number of genes increased.

**Figure 2:**
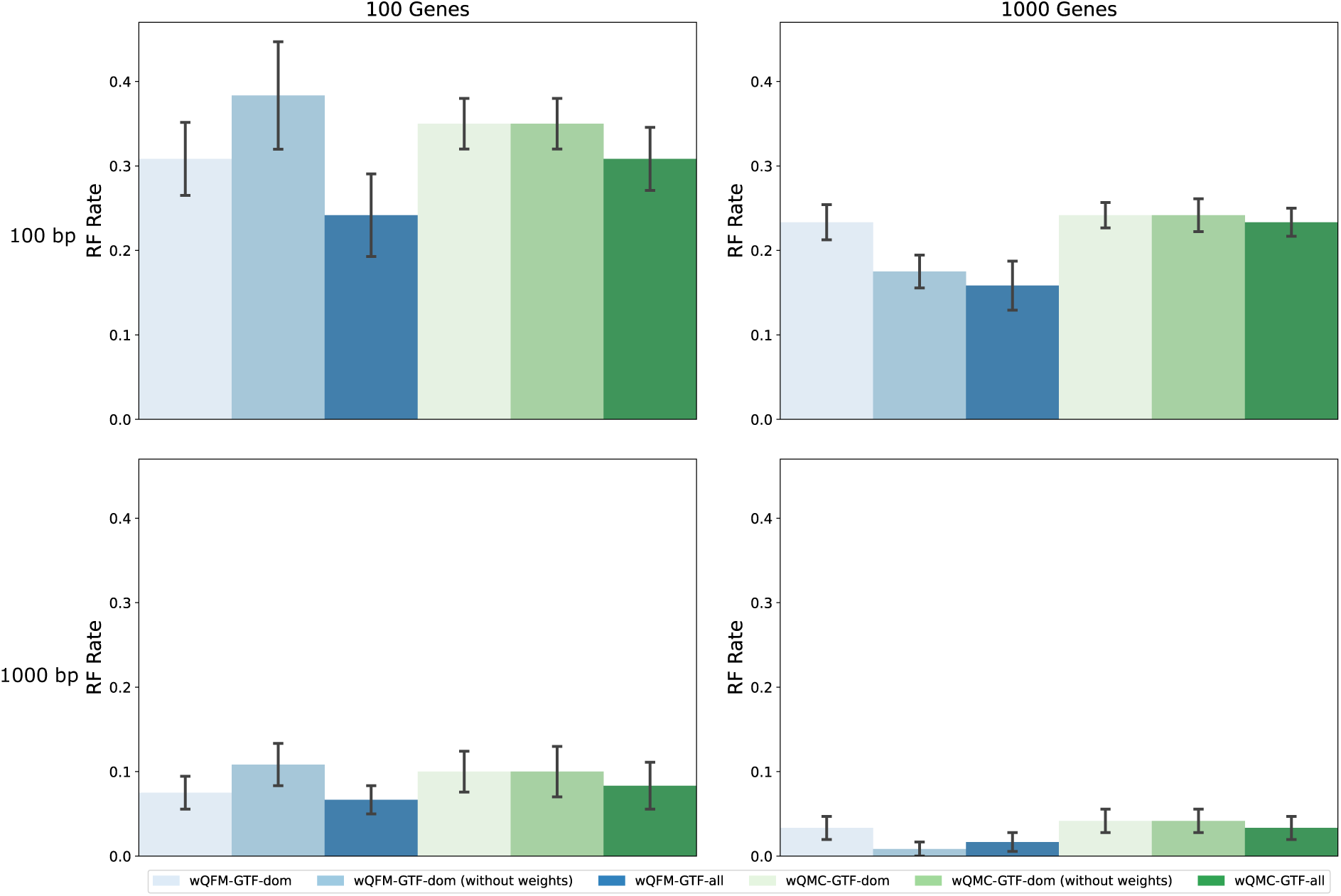
RQ1 (Experiment 1): Results on the 15-taxon dataset. Comparison of performance when using all quartets with weights, as opposed to using only the dominant quartets with and without weights. We show the average RF rates averaged over 10 replicates with standard errors.

We further investigated the performance of these six approaches on the 37-taxon dataset under various model conditions with varying amounts of ILS, numbers of genes, and gene tree estimation errors (Figure 3). Similar trends, as found on 15-taxon datasets, were observed for this dataset as well. All methods showed improved accuracy as we increased the sequence length and number of gene trees and decreased the amount of ILS. wQFM-GTF-all outperformed other methods across all the model conditions. Identical patterns were observed on the 11-taxon dataset (see Supplementary Figure S1). Based on these results, we will consider GTF-all in the subsequent comparisons, while excluding GTF-dom.

**Figure 3:**
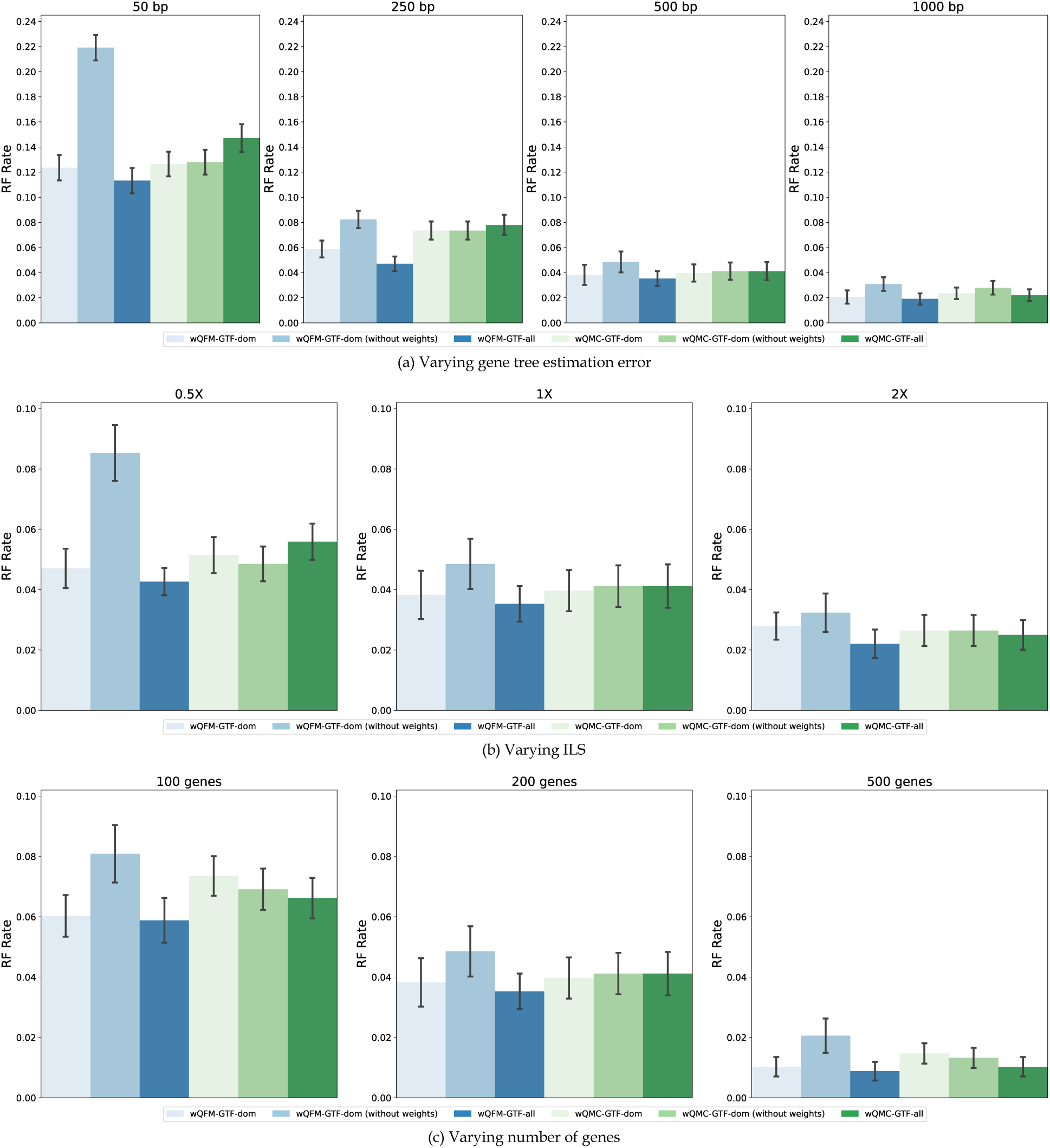
RQ1 (Experiment 1): Results on the 37-taxon dataset. The quartet distributions are analogous to the 15 taxa version. (a) The sequence length was varied from 50 bp to 1000 bp, keeping the ILS fixed at 1X and the number of genes at 200. (b) The ILS was varied from high (0.5X) to low (2X), keeping the sequence length fixed at 500 bp and the number of genes at 200. (c) The number of genes was varied from 100 to 500, keeping the ILS fixed at 1X and the sequence length at 500 bp.

In light of these results, we will only consider the GTF-all strategy in the subsequent comparisons. From now on, we will denote the wQFM and wQMC-based results by wQFM-GTF and wQMC-GTF.

#### Experiment 2: BestML gene trees vs bootstrap/Bayesian distribution of gene trees

In this experiment, we assess the impact of considering a distribution of trees for each gene, rather than a single tree per gene, in weighted quartet-based species tree estimation. Summary methods can utilize either a single maximum likelihood (ML) tree estimate for each gene or a collection of gene trees derived from bootstrap replicates. The former approach, where only the best ML tree for each gene is used, is denoted as BestML. Methods such as wQFM-GTF and wQMC-GTF fall within this category. We seek to understand if considering a distribution of trees for each gene, as methods like BUCKy do, results in improved weighted quartet distribution and eventually more accurate species trees.

We compared the impact of using a distribution of trees from a Bayesian MCMC analysis (Larget and Simon, 1999) with a distribution of trees obtained through nonparametric bootstrapping (Felsenstein, 1985). These two sampling methods are widely used to obtain credible sets of trees and to estimate support values for phylogenies. We used MrBayes and RAxML to generate Bayesian MCMC and nonparametric bootstrap samples of trees, respectively.

Results on the 15-taxon datasets are shown in Figure 4. For the 100bp (shorter sequence length) model condition, methods using distributions of gene trees as input outperformed those using BestML gene trees. Additionally, Bayesian tree distributions resulted in more accurate species trees than ML bootstrapping trees. Notably, wQFM-GTF-MB achieved the highest accuracy across all model conditions, showing substantial improvements over other methods. As the sequence length increased to 1000bp, Bayesian tree distributions continued to produce the best results, whereas the RAxML bootstrapping-based approach performed worse than BestML, regardless of the number of genes.

**Figure 4:**
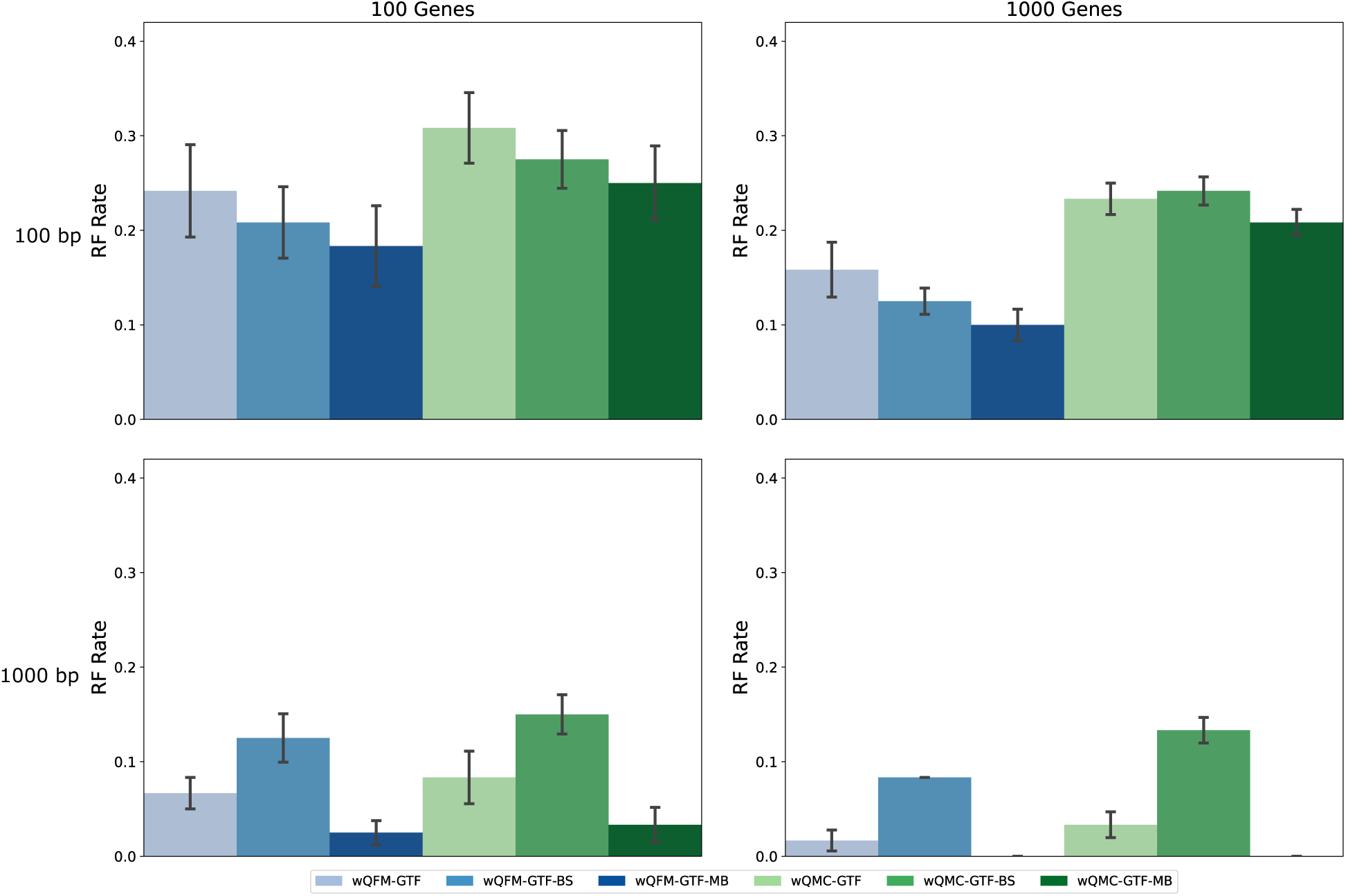
RQ1 (Experiment 2): Results on the 15-taxon dataset. We compare methods utilizing weighted quartets generated from BestML gene trees (GTF), bootstrap distribution of gene trees (GTF-BS), and Bayesian distribution of gene trees (GTF-MB).

Similar trends were observed on the 37-taxon dataset (Figure 5), with the exception that wQFM-GTF-BS (RAxML) consistently produced less accurate trees than wQFM-GTF, even under model conditions with shorter sequence lengths. As observed in other experiments, wQFM produced notably better trees than wQMC. Although wQFM-GTF-MB is better than wQFM-GTF across all model conditions, their difference tends to decrease as we increase the number of genes, the amount of ILS, and the sequence length. The superiority of wQFM-GTF over wQFM-GTF-BS indicates that BestML gene trees tend to produce more accurate quartet tree distributions than non-parametric bootstrapping using RAxML, supporting prior evidence that BestML produces better trees than BS [34]. On the other hand, the enhanced performance of wQFM-GTF-MB over wQFM-GTF suggests that Bayesian gene tree distributions produce more accurate quartet weights than BestML trees.

**Figure 5:**
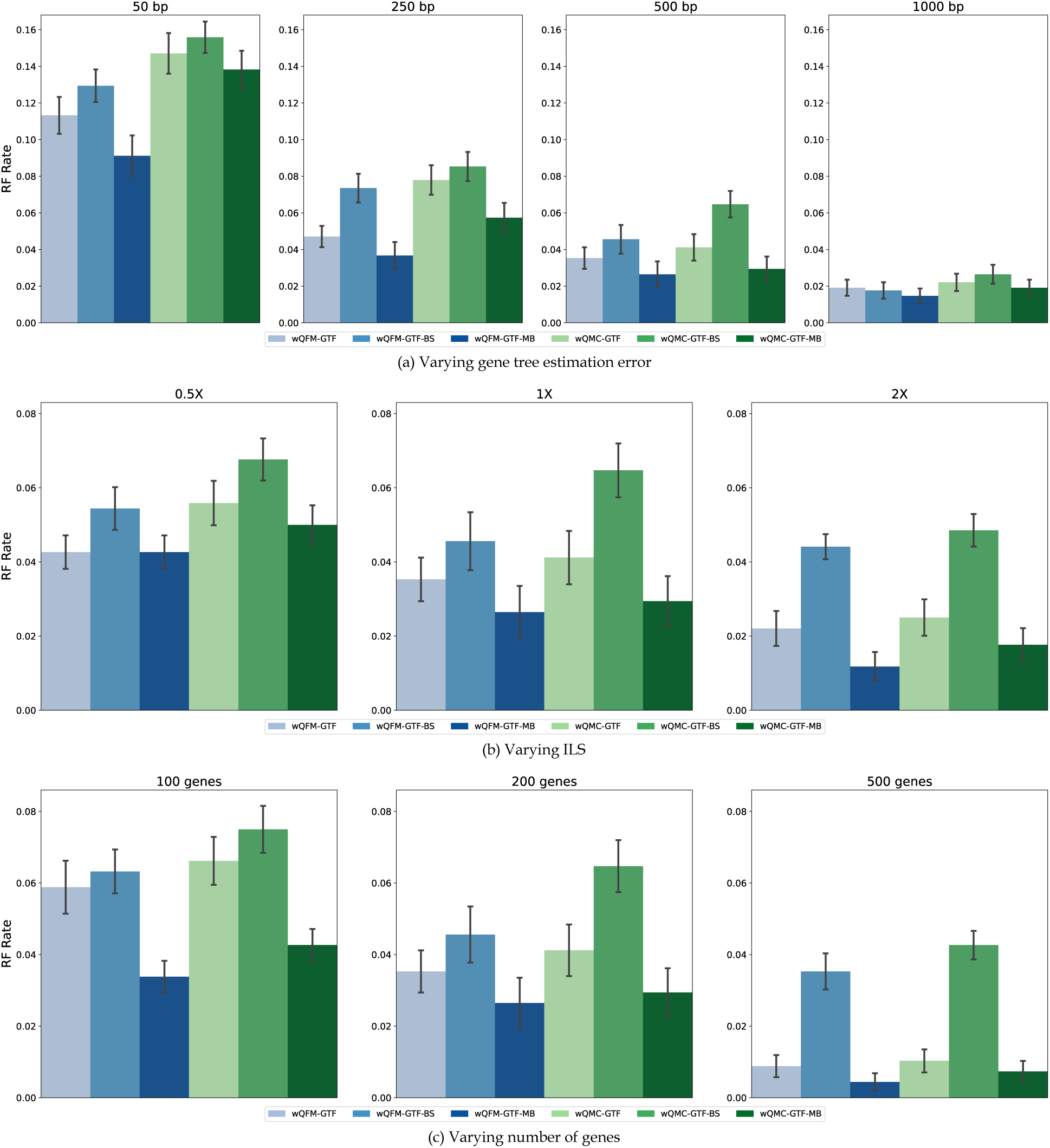
RQ1 (Experiment 2): Results on the 37-taxon dataset. We compare methods utilizing weighted quartets generated from BestML gene trees (GTF), bootstrap distribution of gene trees (GTF-BS), and Bayesian distribution of gene trees (GTF-MB). The model conditions are identical to Figure 3.

To further investigate this, we compared the quartet distributions computed based on, BestML, bootstrapping (BS), and MrBayes (MB) gene trees with those of true gene trees. We measure the divergence between true quartet distributions (computed from true gene trees) and different sets of quartet distributions in estimated gene trees (e.g., BestML, BS, and MB) in terms of the number of dominant quartets that differ between two quartet distributions. Note that, for a set of *n* taxa, there are 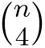 different four-taxa sets, and thus 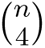 dominant quartets. In Table 2, we show the number of dominant quartets in BestML, BS, and MB sets of gene trees that differ from the dominant quartets in true gene trees. MB trees yield the least number of mismatches in dominant quartets with respect to true dominant quartets. Similarly, the mismatches for BestML are less than BS on the model conditions where wQFM-GTF is better than wQFM-GTF-BS. These are aligned with the relative species tree accuracies of wQFM-GTF-MB, wQFM-GTF-BS, and wQFM-GTF.

**Table 2:**
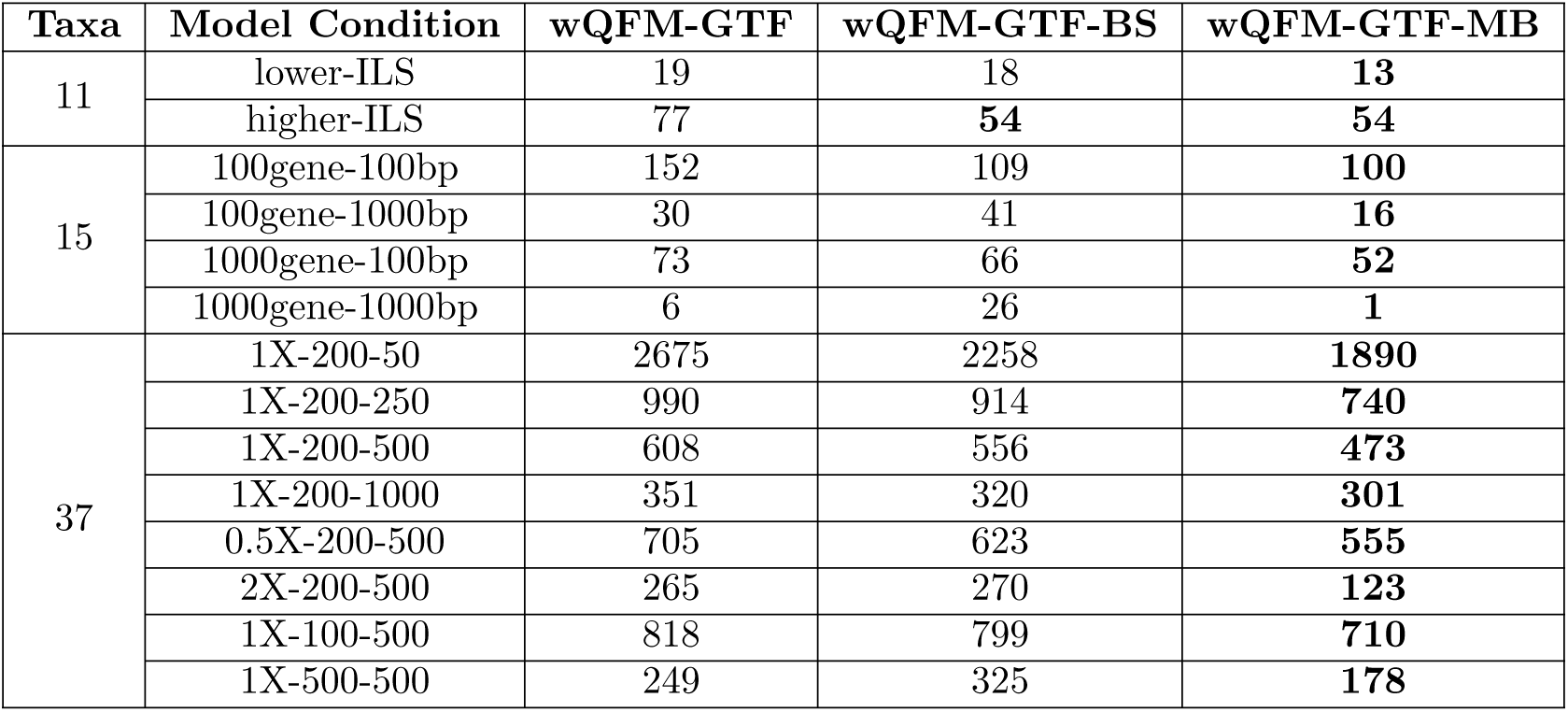
Number of mismatched dominant quartets with respect to the true gene trees for different model conditions averaged over all replicates. The best-performing method (i.e., the lowest number of mismatch) for each model condition has been highlighted in bold.

| Taxa | Model Condition | wQFM-GTF | wQFM-GTF-BS | wQFM-GTF-MB |
| --- | --- | --- | --- | --- |
| 11 | lower-ILS | 19 | 18 | **13** |
|  | higher-ILS | 77 | **54** | **54** |
| 15 | 100gene-100bp | 152 | 109 | **100** |
|  | 100gene-1000bp | 30 | 41 | **16** |
|  | 1000gene-100bp | 73 | 66 | **52** |
|  | 1000gene-1000bp | 6 | 26 | **1** |
| 37 | 1X-200-50 | 2675 | 2258 | **1890** |
|  | 1X-200-250 | 990 | 914 | **740** |
|  | 1X-200-500 | 608 | 556 | **473** |
|  | 1X-200-1000 | 351 | 320 | **301** |
|  | 0.5X-200-500 | 705 | 623 | **555** |
|  | 2X-200-500 | 265 | 270 | **123** |
|  | 1X-100-500 | 818 | 799 | **710** |
|  | 1X-500-500 | 249 | 325 | **178** |

This supports prior evidence that BestML tends to produce better trees than BS. But we did not explore MLBS – future work.

In the 11-taxon dataset, the difference between the methods is less prominent (Figure S2). On higher ILS, the bestML trees perform poorly. But other than that, all methods obtain comparable results. There is not much distinction between wQFM and wQMC either, unlike the previous two datasets.

#### Experiment 3: Utilizing the unweighted and weighted quartets generated by SVDquar-tets

SVDquartets generates dominant quartets based on the SVD-score and then amalgamates them using QFM or QMC in an unweighted setting. It can also generate weighted quartets in two settings: exponential and reciprocal. These weighted quartets can be used with wQFM or wQMC to estimate species trees. However, although SVDquartets is widely studied and has been compared to coalescent-based methods [35], to the best of our knowledge, the performance of the weighted settings in SVDquartets has not been previously explored. In this experiment, we compared standard SVDquartets with wQFM and wQMC when they were run on the weighted quartets generated by SVDquartets (i.e., wQFM-SVD-Exp, wQFM-SVD-Rec).

Comparisons of these methods on 15-and 37-taxon datasets are shown in Figures 6 and 7. Results on the 11-taxon dataset are presented in Supplementary Material (Figure S3). SVDquartets, wQFM-SVD-Rec, and wQMC-SVD-Exp showed comparable performance, but wQFM-SVD-Rec was better than SVDquartets on some conditions (e.g., 1000-gt model condition in the 15-taxon dataset). wQFM-SVD-Exp and wQMC-SVD-Rec performed worse than others in many model conditions.

**Figure 6:**
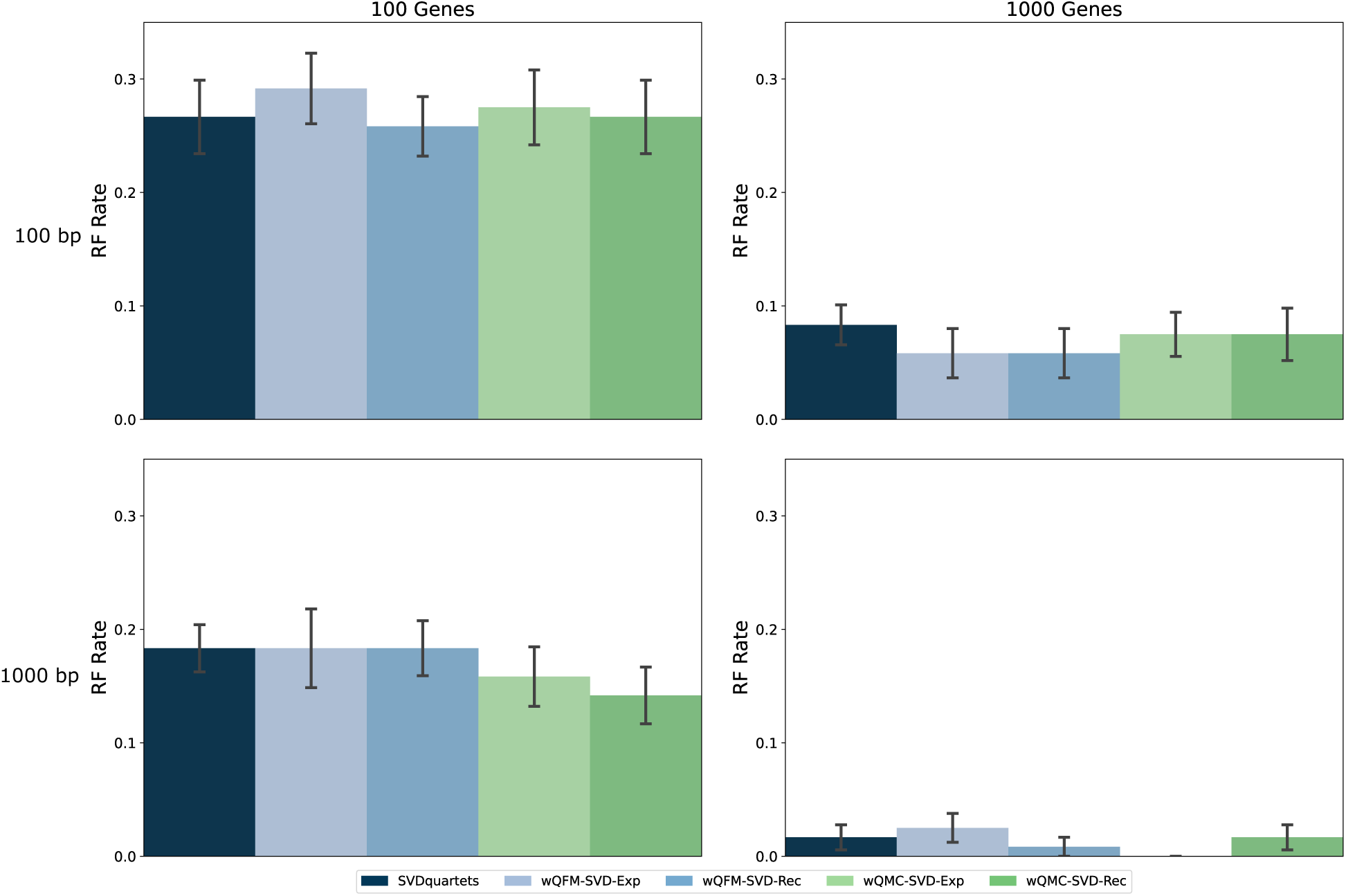
RQ1 (Experiment 3): Results on the 15-taxon dataset. Performance compar-ison between unweighted and weighted (both exponential and reciprocal) quartets generated by SVDquartets. The unweighted quartets are amalgamated by QFM, whereas the weighted ones are processed by both wQFM and wQMC.

**Figure 7:**
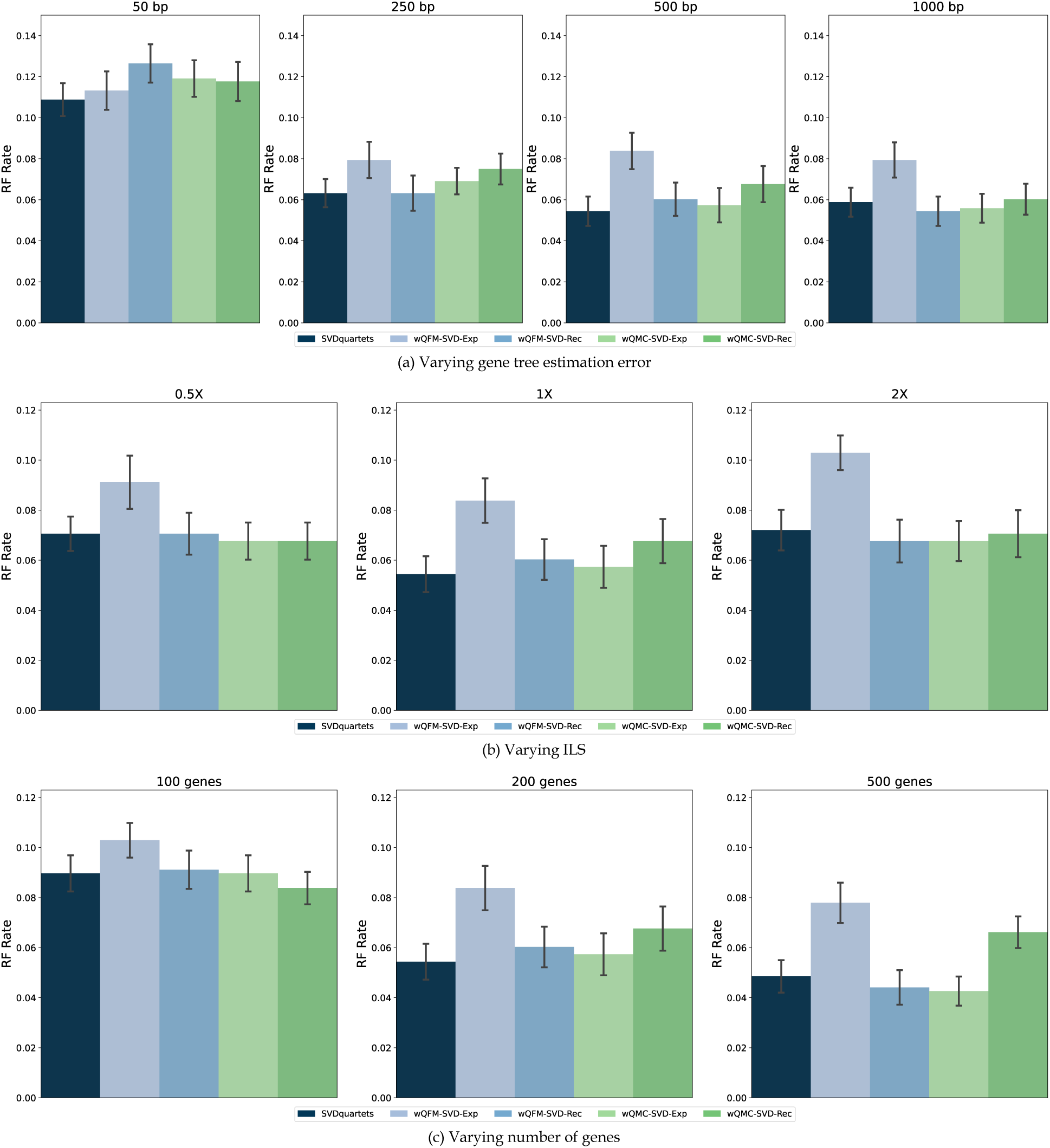
RQ1 (Experiment 3): Results on the 37-taxon dataset. Performance compar-ison between unweighted and weighted (both exponential and reciprocal) quartets generated by SVDquartets. The unweighted quartets are amalgamated by QFM, whereas the weighted ones are processed by both wQFM and wQMC. The settings are identical to Figure 3.

Interestingly, contrary to the expected trends, the trees estimated by SVDquartets and those based on the weighted quartets generated by SVDquartets do not show notable improvements with increased sequence lengths or decreased levels of ILS. As shown in Figure 7(a), expected improvements were observed when the sequence length increased from 50bp to 250bp; however, no further improvement was noted beyond 250bp. Similarly, no improvements were observed with decreased levels of ILS (Figure 7(b)), and surprisingly, it produced the worst results under the lowest-ILS (2X) model condition. However, with an increase in the number of genes, the results improved as expected. Overall, wQFM-SVD-Rec performs better than other variants of SVDquartets explored in this particular experiment.

#### Experiment 4: Utilizing the weighted quartets generated by BUCKy and maximum likelihood (ML) based methods

We investigated different variations of Maximum Likelihood (ML) and BUCKy-based methods and even some form of combinations between them.

We used MrBayes and RAxML to generate gene tree distributions as input to BUCKy (denoted by BUCKy-MB and BUCKy-RAxML, respectively). BUCKy computes the concordance factors of each quartet tree. The concordance factor of a quartet is the proportion of gene trees that truly have that quartet [26]. Thus, these concordance factors measure the genomic support of each quartet. In this experiment, in addition to evaluating the BUCKy-pop tree, we used the CFs of the quartets generated by BUCKy as weights and amalgamated them using wQFM and wQMC to estimate the species tree.

The results on the 37-taxon datasets are presented in Figure 9. BUCKy-MB and wQFM-GTF achieved the highest accuracy. Notably, BUCKy-MB significantly outperformed BUCKy-RAxML, demonstrating the superiority of gene trees estimated by Bayesian techniques (MrBayes) over those derived from ML-based techniques [36, 37]. This further supports our findings in Experiment 2. However, wQFM-BUCKy-MB and wQFM-BUCKy-RAxML performed worse than other methods, suggesting that using CFs as weights for quartets and estimating species trees from these weighted quartets using quartet amalgamation techniques may not be effective. Another key observation is that, unlike SVDquartets, increasing sequence length, and thereby reducing gene tree estimation error, significantly reduces the error rate of BUCKy-estimated trees.

Similar trends were observed on 15-and 11-taxon datasets (see Figures 8 and S4). In this experiment, we also investigated the efficacy of the weighted quartets generated by RAxML in species tree estimation. We ran wQFM on the quartets weighted based on the likelihood scores computed by RAxML. This approach (wQFM-RAxML) performed poorly (see Supplementary Figure S5).

**Figure 8:**
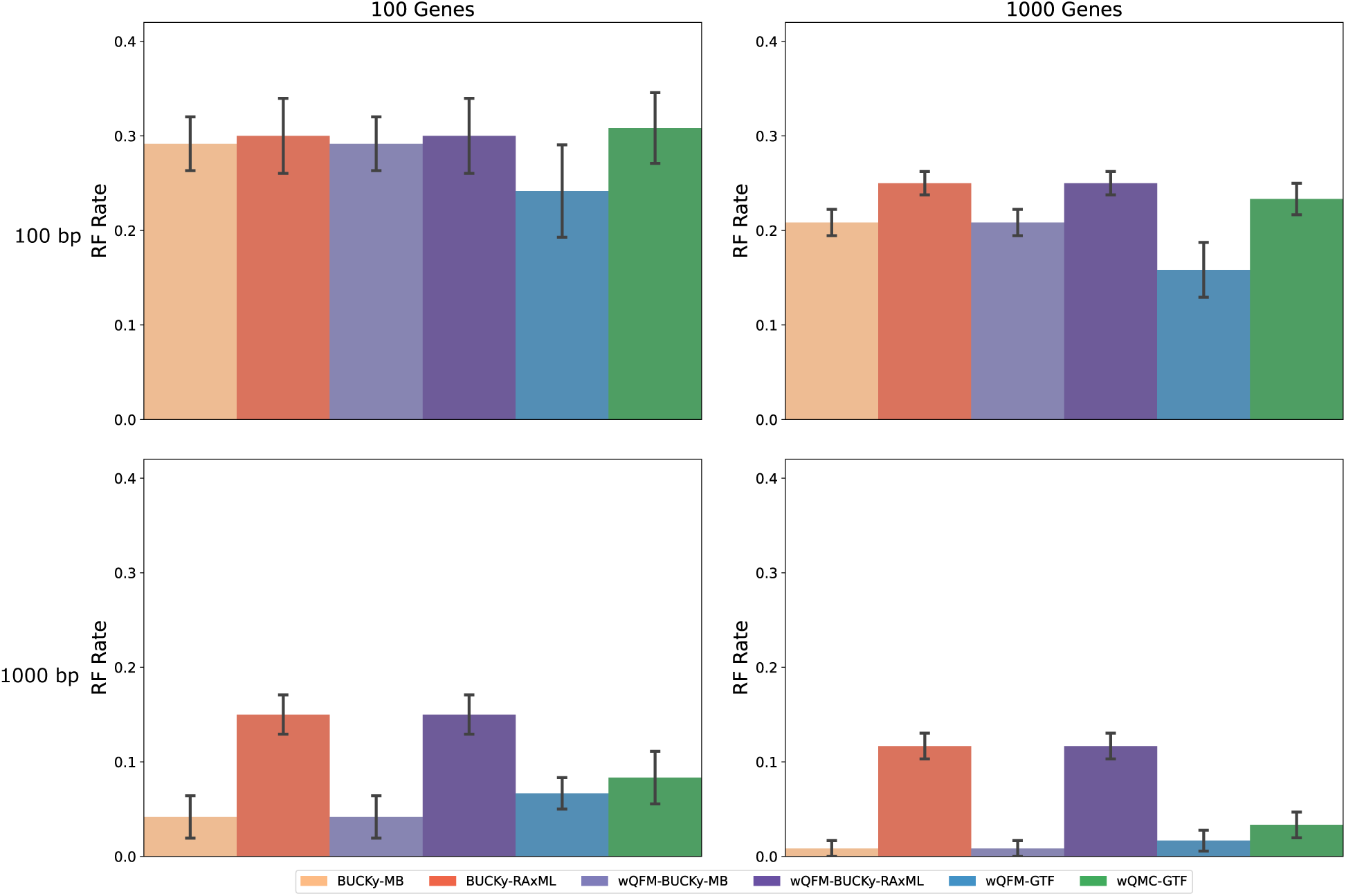
RQ1 (Experiment 4): Results on the 15-taxon dataset. Comparison of various BUCKy-based and ML-based methods. We also included wQFM-GTF and wQMC-GTF.

**Figure 9:**
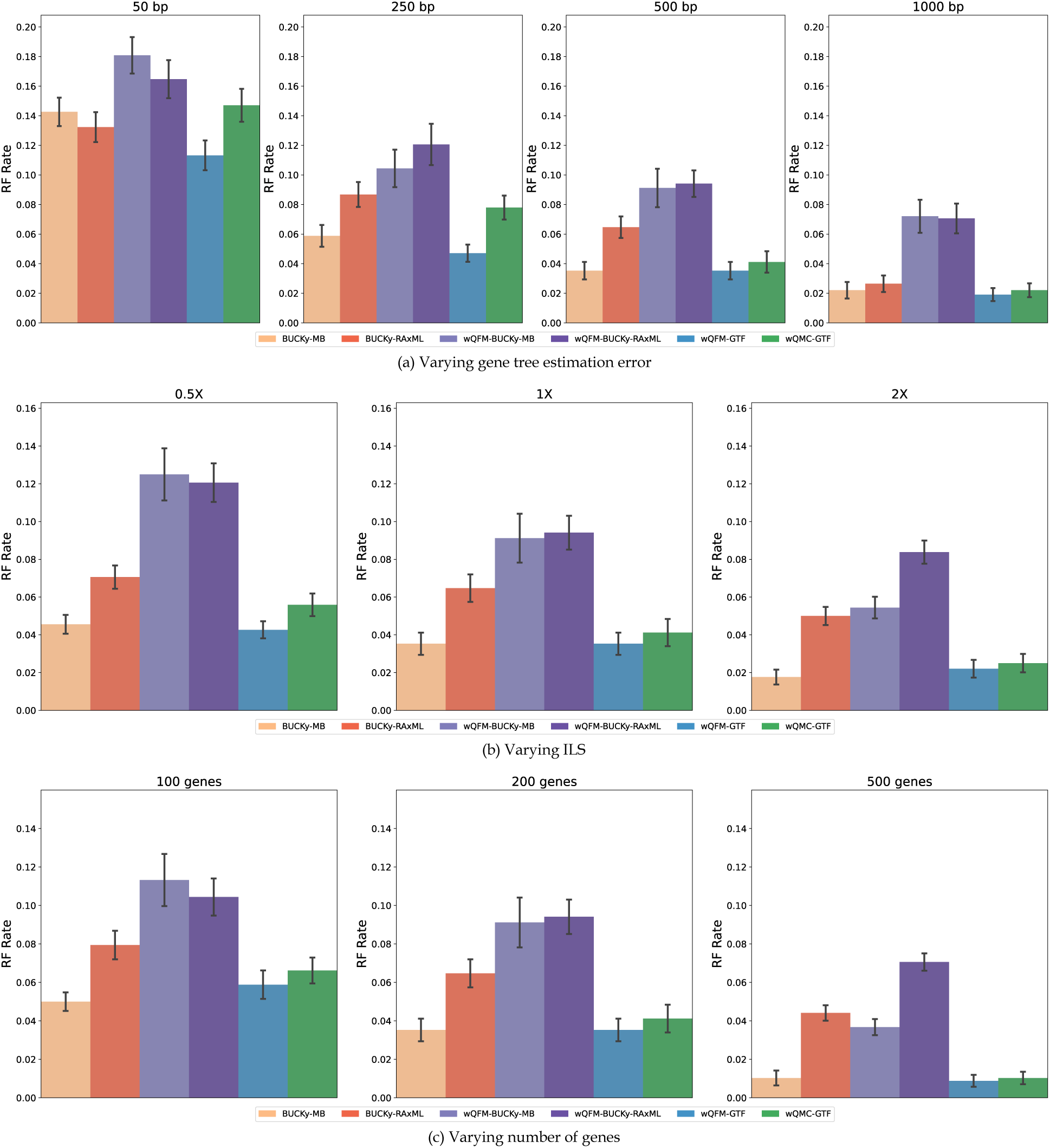
RQ1 (Experiment 4): Results on the 37-taxon dataset. Comparison of various BUCKy-based and ML-based methods. We also included wQFM-GTF and wQMC-GTF. The model conditions are identical to Figure 3.

This particular set of analyses presents us with a method that is clearly superior to others, wQFM-GTF-MB.

### 3.2 RQ2: Investigating the relative performance of different weighted quartet amalgamation techniques

wQFM and wQMC are two leading weighted quartet amalgamation techniques. Prior studies [9, 38] demonstrated that wQFM-GTF substantially outperforms wQMC-GTF. In addition to GTF, this study evaluates both wQFM and wQMC using a broad range of weighted quartet generation methods. Our extensive evaluation (as discussed in Experiments 1-4 under RQ1) supports previous findings on the relative performance of wQFM and wQMC, further establishing the superiority of wQFM. It consistently outperformed wQMC across various model conditions, with many differences being statistically significant. As a result, wQMC is excluded from further experiments in RQ3-RQ5. Additionally, Table S1 shows that trees estimated by wQFM attain “closer” quartet scores to the true tree compared to wQMC, proving that wQFM is more resistant to gene tree estimation errors. These results have been explained in more detail in Section 3.4.

### 3.3 RQ3: Relative performance of different quartet-based summary methods

We compared the best weighted quartet amalgamation approaches identified in RQ1—wQFM-GTF, wQFM-GTF-MB, and wQFM-SVD-Rec—with the leading quartet-based summary methods ASTRAL, BUCKy, and SVDquartets. Both ASTRAL and its weighted variant (wASTRAL) were analyzed, with wASTRAL utilizing weights derived from branch supports. Our experiments, particularly Experiment 2 in RQ1, demonstrated that leveraging a distribution of trees for each gene, rather than a single BestML tree per gene, enhances tree accuracy. Consequently, we ran wASTRAL on BestML gene trees with supports estimated from both non-parametric bootstrapping (denoted as wASTRAL) and Bayesian MCMC sampling (wASTRAL-MB). Thus, we used the same distributions of trees as input for both wQFM-GTF-MB and wASTRAL-MB, ensuring a fair comparison. These tree distributions were utilized to generate weighted quartets for wQFM and to draw branch support on the gene trees used as input for ASTRAL.

Results on 15-and 37-taxon datasets are shown in Figures 10 and 11. The general patterns were consistent with our expectations: for all methods, the species tree estimation accuracy was improved by increasing the number of genes and sequence lengths (i.e., decreasing the gene tree estimation error), but was reduced by increasing the amount of gene tree discordance (i.e., the amount of ILS).

**Figure 10:**
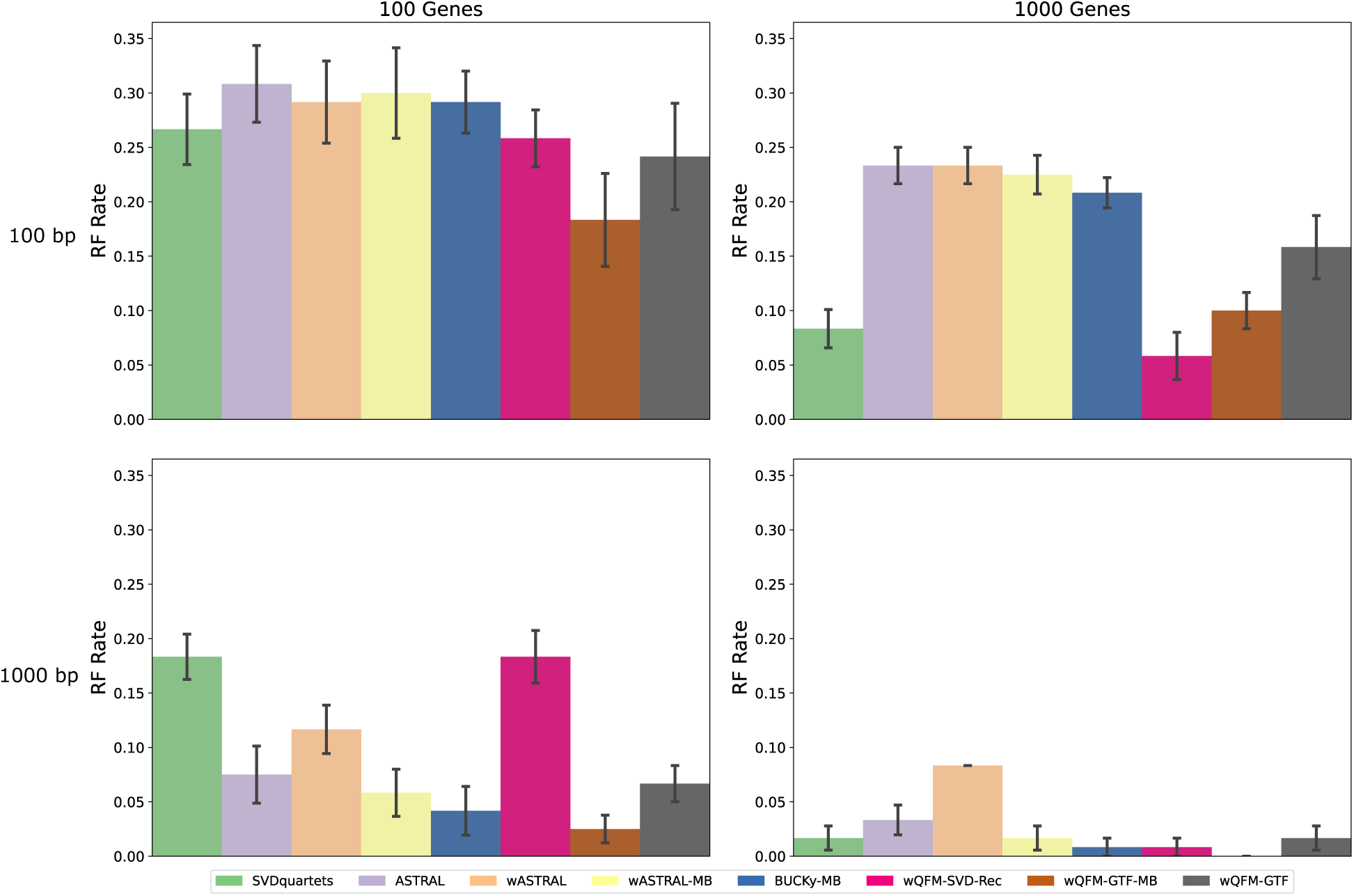
RQ3: Results on the 15-taxon dataset. We compare the best methods from previ-ous experiments: wQFM-GTF, wQFM-GTF-MB, wQFM-SVD-Rec with ASTRAL, BUCKy, and SVDquartets. ASTRAL’s weighted counterpart, configured to utilize branch supports as weights, was also analyzed. Branch supports were estimated from both non-parametric RAxML bootstrap-ping (wASTRAL) and Bayesian MCMC sampling (wASTRAL-MB).

**Figure 11:**
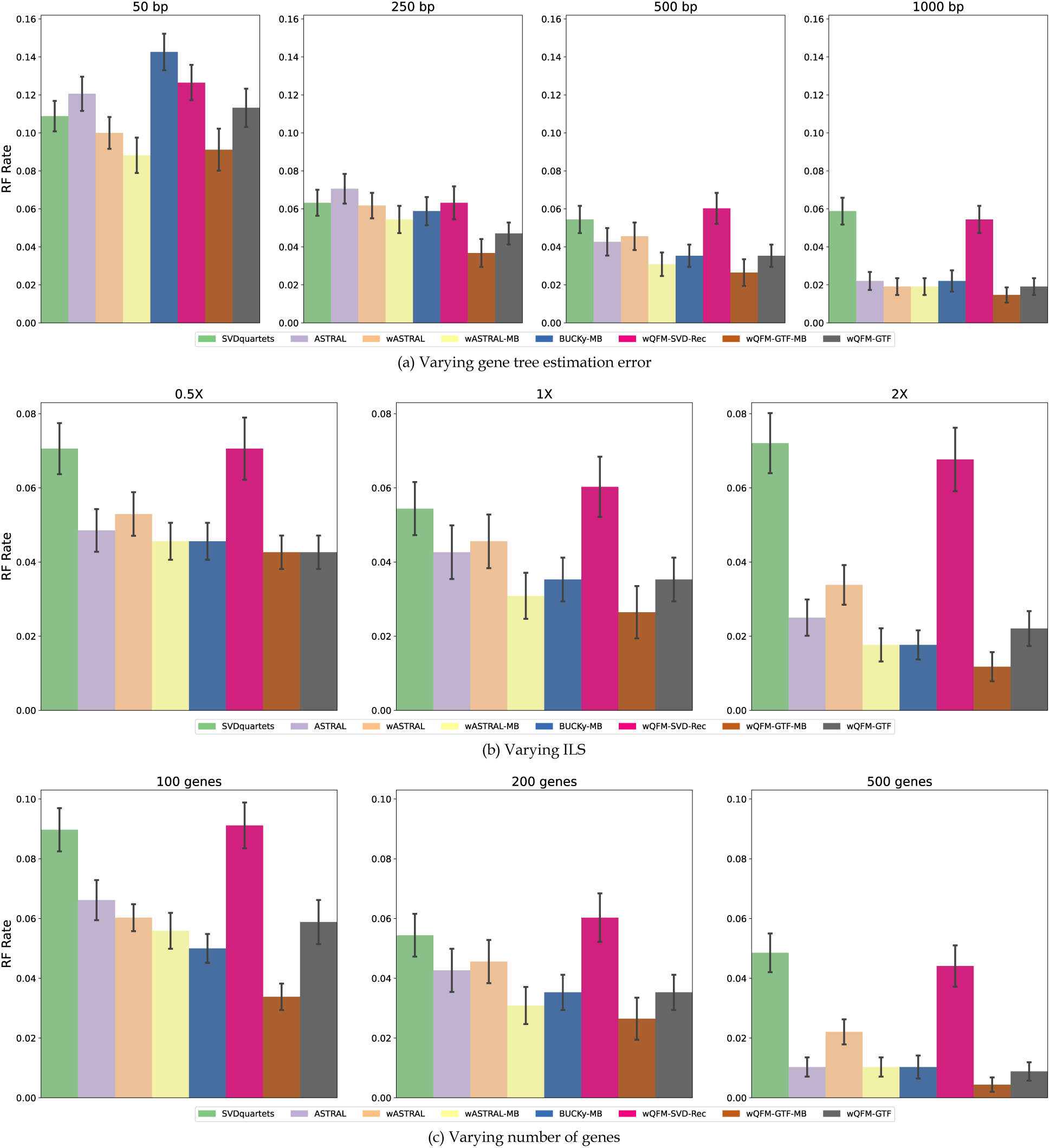
RQ3: Results on the 37-taxon dataset. We compare the best methods from previous experiments: wQFM-GTF, wQFM-GTF-MB, wQFM-SVD-Rec with ASTRAL, BUCKy, and SVDquartets. ASTRAL’s weighted counterpart, configured to utilize branch supports as weights, was also analyzed. Branch supports were estimated from both non-parametric RAxML bootstrapping (wASTRAL) and Bayesian MCMC sampling (wASTRAL-MB). The settings are identical to Figure 3.

Next, we compared the relative performance of different methods. The most significant obser-vation is that wQFM-GTF-MB achieved the best overall performance, substantially outperforming both ASTRAL and wASTRAL. Utilizing MB-estimated tree distributions improves the perfor-mance of both wQFM and ASTRAL. Consistent with the trend observed in Experiment 2 of RQ1, where wQFM-GTF-MB significantly outperformed wQFM-GTF, we found that wASTRAL-RAxML and wASTRAL-MB also outperformed ASTRAL. However, irrespective of the set of trees used, wQFM consistently matched or exceeded the accuracy of its ASTRAL counterparts. Addi-tionally, BUCKy-MB frequently produced more accurate trees than ASTRAL. wQFM-GTF-MB, BUCKy-MB, and wQFM-GTF yield the second-best accuracy, which either matched or outper-formed ASTRAL.

To further investigate the relative performance of these methods and to understand why wQFM performs better than ASTRAL, we computed the quartet scores of the estimated species trees as well as the true species tree with respect to both estimated and true gene trees. These quartet scores, further analyzed in Section 3.4, indicate that although ASTRAL-estimated trees have lower overall tree accuracy compared to wQFM, ASTRAL achieves higher quartet scores relative to the estimated gene trees used as input. Interestingly, however, wQFM achieves higher quartet scores than ASTRAL when these scores are computed with respect to the true gene trees. Furthermore, regardless of whether true or estimated gene trees are used to compute the quartet scores, the quartet scores of wQFM-estimated trees are closer to the true quartet score (i.e., the quartet score of the true species tree) than those of ASTRAL-estimated trees. That means ASTRAL tends to “overestimate” the quartet scores on estimated gene trees and “underestimate” them when scores are computed based on true gene trees. These explain why wQFM is generally more accurate than ASTRAL and also suggest that wQFM is less susceptible to gene tree estimation errors compared to ASTRAL.

The 11-taxon dataset reveals almost identical patterns to the 37-taxon dataset (see Figure S6 in the supplementary material). The only exception is that ASTRAL performs better than BUCKy-MB.

### 3.4 RQ4: Are the quartet scores predictive of species tree accuracy?

In this research question, we examine the relationship between the quartet scores and the RF rates of the most accurate species tree estimation approaches, namely wQFM-GTF, wQFM-GTF-MB, and ASTRAL. All these methods generate the species trees by maximizing the quartet score. The statistical consistency of estimating species trees by maximizing the quartet score criterion holds when a sufficiently large number of true gene trees (without estimation error) are available. However, in practice, the number of genes is limited, and estimated gene trees often contain errors. Consequently, the trees that optimize the quartet score may not correspond to the true species tree. As a result, quartet-based methods may “overshoot” the quartet score, returning trees with higher scores than the true quartet score, particularly when we have a limited number of estimated gene trees (with estimation errors) [38, 39].

We examined the quartet scores of each estimated species tree as well as the true tree with respect to two sets of gene trees: the true ones and the estimated ones. We vary the number of genes and the gene tree estimation error (controlled by gene sequence lengths). As shown in Table 3, the quartet score of the true species tree is the highest with respect to true gene trees (which is aligned with the statistical consistency of quartet score). However, with respect to the estimated gene trees (with error), the true species tree may have lower quartet scores. Thus, in the presence of gene tree estimation error and limited numbers of genes, quartet-based methods may “overshoot” the quartet score as they return trees with higher quartet scores than the true quartet score. Similarly, the most accurate method wQFM-GTF-MB yields higher quartet scores than wQFM-GTF and ASTRAL with respect to true gene trees, but ASTRAL and wQFM-GTF produce higher quartet scores than wQFM-GTF-MB (although they are less accurate than wQFM-GTF-MB) when the scores are computed with respect to estimated gene trees. Thus, the quartet score wQFM-GTF-MB is closer to the true quartet score than wQFM-GTF, wQFM-GTF-BS, and ASTRAL, regardless of the gene tree set used to compute the quartet score.

**Table 3:**
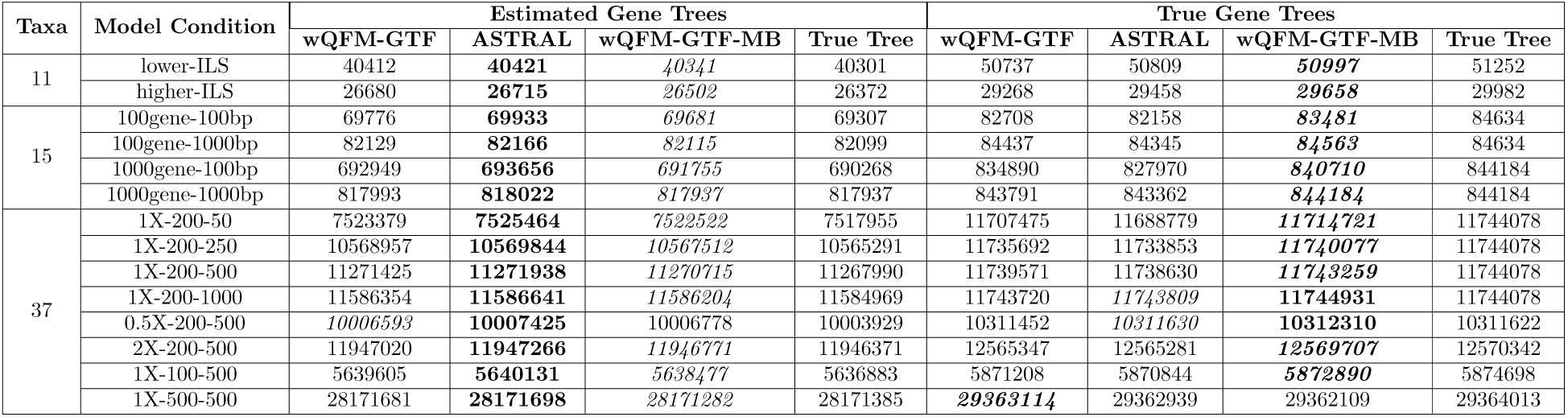
Quartet scores of selected methods for estimated and true gene trees in different model conditions. The highest scores are shown in boldface, and the scores closest to the true score have been italicized.

We visualize the quartet scores for the 15-taxon dataset in Figure 12 to better understand this trend. This figure suggests that the expected trend that quartet scores decrease with increasing RF rates holds when the quartet scores are computed based on true gene trees (blue lines), and the true species tree has the highest quartet score. Interestingly, however, when the quartet scores are based on estimated gene trees (orange lines), the estimated trees may achieve higher quartet scores than the true species tree – meaning that quartet-based methods may “overestimate” quartet scores under challenging model conditions with limited numbers of estimated gene trees (with estimation error). However, regardless of the choice of input gene trees, more accurate methods tend to produce quartet scores that are “closer” to true quartet scores. As demonstrated in Figure 12, quartet scores of the most accurate method wQFM-GTF-MB represented by triangles are closer to true quartet scores (filled rectangles) than other methods in both blue and orange lines. As expected, when we increase the number of genes or decrease the gene tree estimation error (by increasing sequence length)–thereby making the model conditions favorable for statistically consistent criteria like quartet score maximization–the issue with overshooting the quartet scores decreases, evident from the gentler slopes.

**Figure 12:**
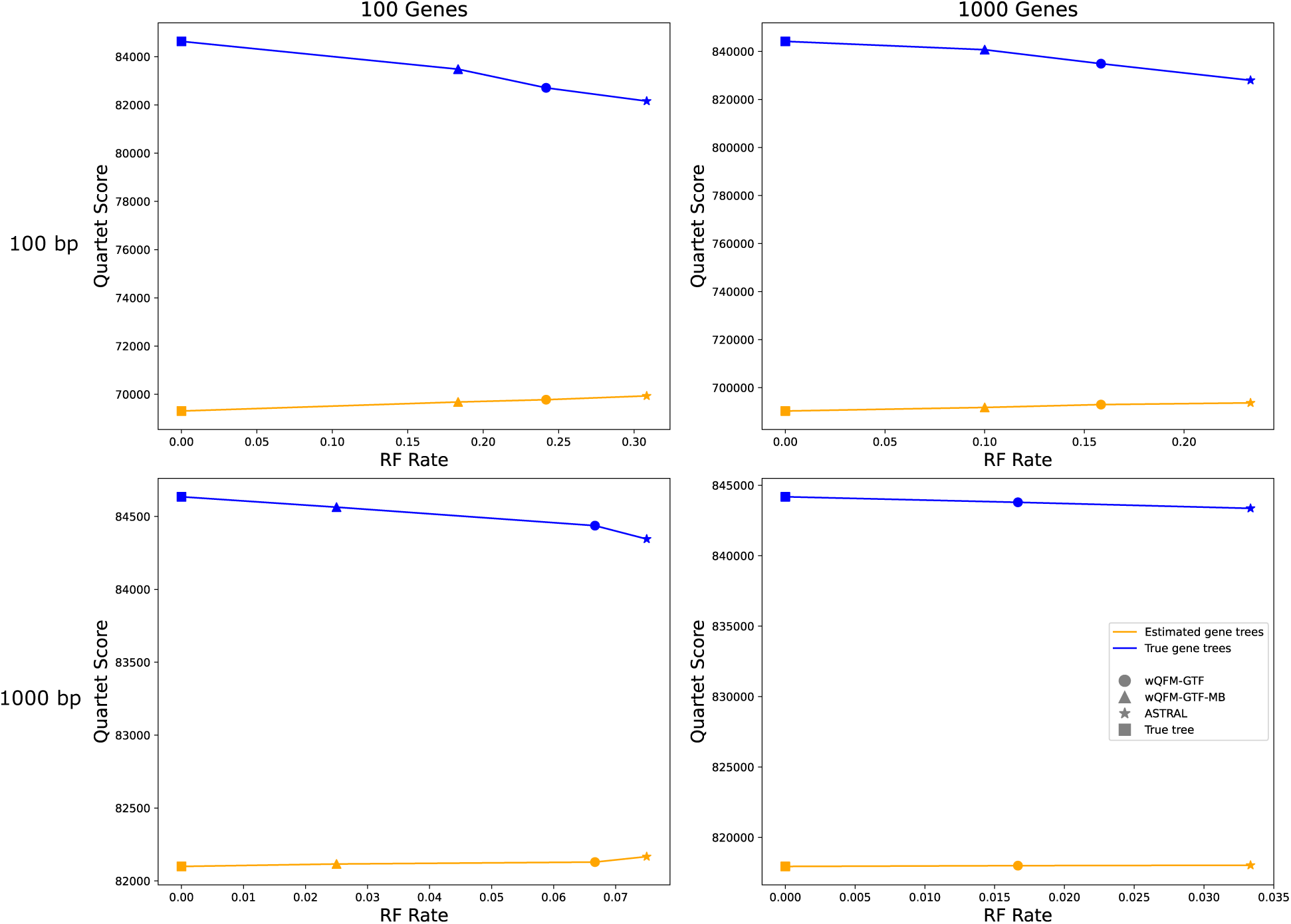
RQ4: Results on the 15-taxon dataset. We show the relation between quartet score and RF rate with respect to both true gene trees and estimated gene trees for different methods (wQFM-GTF, wQFM-GTF-MB, ASTRAL).

Additionally, as discussed in Section 3.2, we compare the quartet scores for wQFM and wQMC in Table S1. Consistent with earlier findings, wQFM, the superior method, exhibits smaller deviations from the true quartet scores compared to wQMC. When using estimated gene trees, wQMC tends to overestimate the quartet scores, whereas for true gene trees, it underestimates them, yielding scores lower than the actual quartet score.

### 3.5 RQ5: Performance on real biological dataset

#### 3.5.1 Mammalian dataset

Song *et al.* analyzed a dataset containing 447 genes across 37 mammals using MP-EST [40] and concatenation using maximum likelihood [32]. We re-analyzed the mammalian dataset from [32] after removing 21 mislabeled genes (confirmed by the authors), and two other outlier genes. The placement of bats (*Myotis lucifugus* and *Pteropus vampyrus*) and tree shrews (*Tupaia belangeri*) were two of the questions of greatest interest, and alternative relationships have previously been reported [41–44].

The trees produced by ASTRAL, BUCKy, and wQFM with different types of tree distributions are identical to each other (see Fig. 13). This tree placed tree shrews (*Tupaia belangeri*) as sister to Glires with high support, which is consistent to the CA-ML analyses (reported in [32]), and bats have been placed as sister to the clade containing Cetartiodactyla, Carnivora, and Perissodactyla (which is consistent to the MP-EST analyses [34]). The SVDquartets-estimated tree is identical to this tree except for the position of tree shrews. SVDquartets supports an alternative relationship (albeit with very low support), which has also been observed [8,34], that placed tree shrews as sister to Glires. With respect to the position of bats, all the trees reconstructed in this study agree with MP-EST, which placed bats as sisters to the (Cetartiodactyla, (Perissodactyla, Carnivora)) clade. The placement of tree shrews and bats is of substantial debate, and so the differential placement is of considerable interest in mammalian systematics.

**Figure 13:**
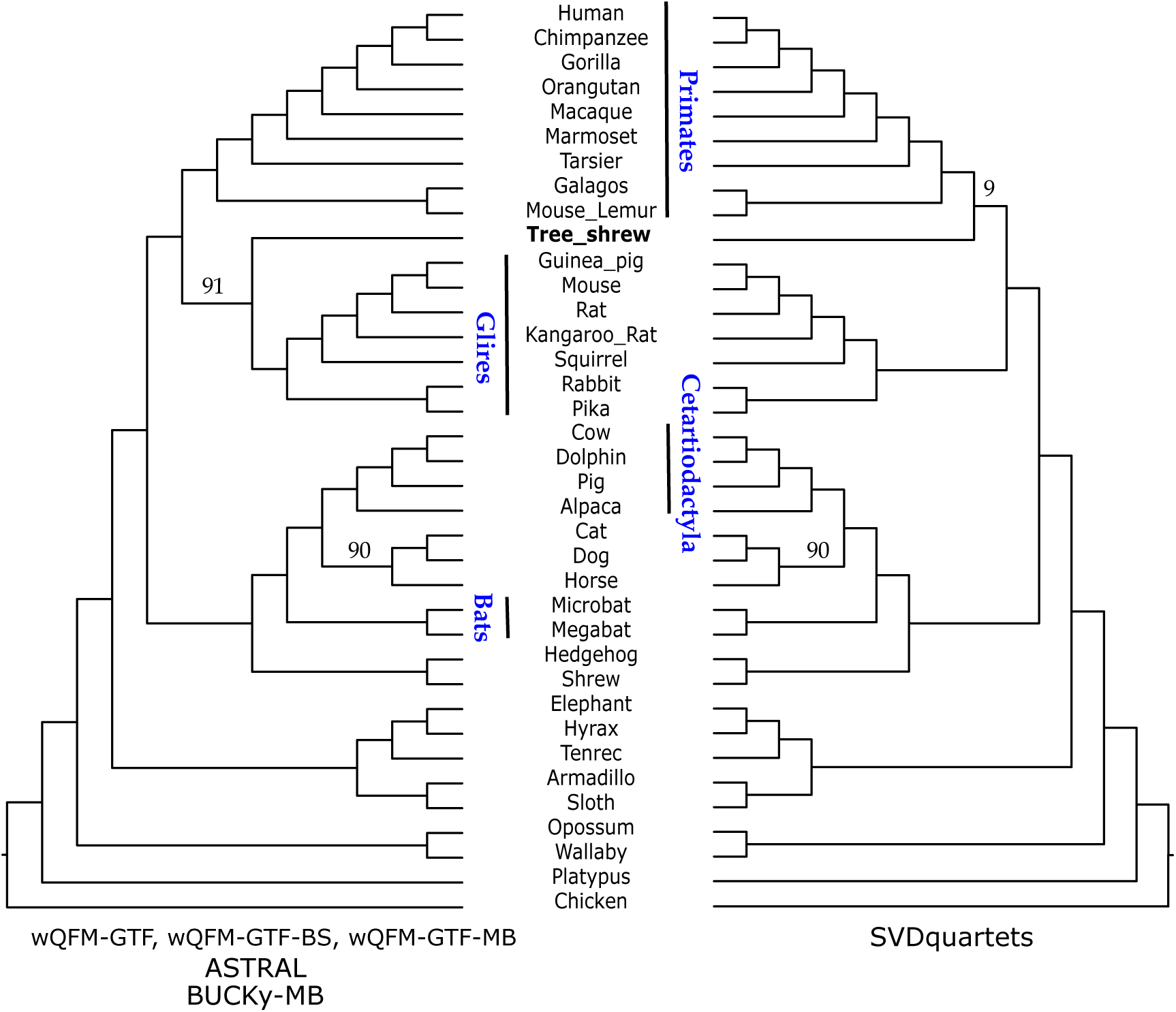
Analysis of the mammalian dataset. We show the trees estimated by different meth-ods, e.g., ASTRAL, BUCKy, SVDquartets, and wQFM with different types of quartet distributions. Branch supports are computed based on quartet-based local posterior probability (multiplied by 100). All BS values are 100% except where noted.

#### 3.5.2 Avian dataset

We reanalyzed the avian biological dataset from Jarvis et al. [3], which comprises 14,446 genes across 48 taxa, including exons, introns, and ultra-conserved elements (UCEs). This dataset poses significant challenges due to high levels of gene tree discordance, likely driven by rapid radiation events in the evolutionary history of these species. Mahbub *et al.* [9] provided a comparison between the reference tree (MP-EST with statistical binning), ASRAL, wQFM, and wQMC. In this study, we extend this analysis to evaluate the performance of wQFM-GTF-BS and SVDquartets (see Figure. 14). Due to the computational demands of generating Bayesian distributions for each of the 14K gene trees, we were unable to include wQFM-GTF-MB and BUCKy-MB in our current analysis.

**Figure 14:**
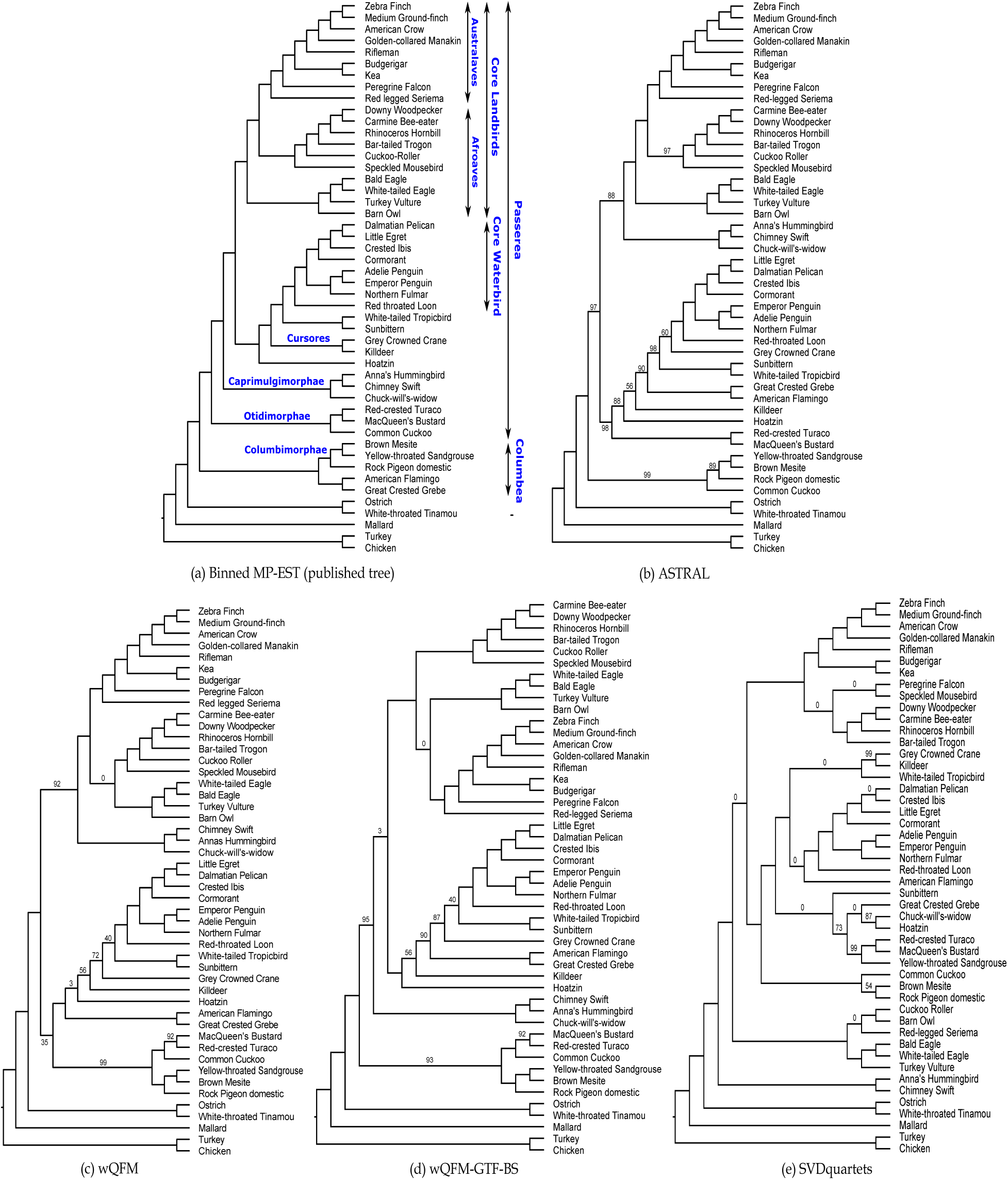
Analysis of the avian dataset. We show the trees estimated by different methods, e.g., ASTRAL, SVDquartets, wQFM, and wQFM-GTF-BS. Branch supports are computed based on quartet-based local posterior probability (multiplied by 100). All BS values are 100% except where noted.

ASTRAL, wQFM, and wQFM-GTF-BS produced reasonably good phylogenies, although neither successfully reconstructed several sub-groups that have been consistently supported in the avian phylogenomics project and other studies [9]. Notably, both the wQFM-estimated trees showed greater congruence with the MP-EST tree (derived from binned gene trees) than the ASTRAL-estimated tree. Surprisingly, SVDquartets yielded a significantly poorer tree, failing to recover many of the established sub-groups. All three of wQFM, wQFM-GTF-BS, and ASTRAL accurately reconstructed the well-established Australaves clade, containing passeriformes, parrots, falcons, and seriemas. But SVDquartets misplaced both Seriema (with 100% support) and Falcon, thus failing to reconstruct Australaves.

All four methods failed to recover Columbea. While wQFM, wQFM-GTF-BS, and ASTRAL were able to recover the constituent clades Columbimorphae (mesite, sandgrouse, and pigeon) and Phoenicopterimorphae (flamingo and grebe), SVDquartets failed to recover any of the two. Addi-tionally, SVDquartets was unable to resolve other key clades, such as Otidimorphae, Caprimulgimorphae, and Afroaves. Although it successfully recovered the Core Waterbird clade, the relationships within this group were not well-resolved. Interestingly, it managed to recover Cursores (crane and killdeer) with high support, a clade that all other methods failed to reconstruct.

## 4 Conclusions

Highly accurate and scalable species tree inference methods, which summarize independently inferred gene trees to obtain a species tree, are sensitive to unavoidable errors introduced in the gene tree estimation step. This dilemma has sparked significant debate about the advantages of concatenation versus summary methods and has presented practical challenges to the broader adoption of summary methods over concatenation. Co-estimation of gene trees and species trees [45–47] is perhaps the most accurate approach to dealing with such noise [28, 48]. However, despite some progress [49], these methods are extremely computationally intensive and do not scale to even moderately large numbers of species [14, 28, 50]. Therefore, summary methods are comparatively more feasible for use on genome-scale datasets.

Quartet-based summary methods, including ASTRAL and quartet amalgamation-based methods wQFM and wQMC, have emerged as a highly accurate and scalable approach for estimating species trees from multi-locus data. Prior studies suggest that weighting schemes may improve the performance of quartet-based methods. In this study, we proposed and examined various ways of generating weights for the quartets and their impact in species tree inferences. Moreover, we report on an extensive evaluation study, the relative performance of the leading quartet-based methods.

This study shows various important trends, which we summarize. The first observation is that the species trees estimated by quartet amalgamation-based methods were impacted by the choice of input quartet tree distribution (e.g., dominant or all quartets) and that using all quartets with relative weights may better capture the gene tree uncertainty and produced more accurate species tree topologies, especially in the presence of gene tree estimation errors. This study also shows that wQFM produces substantially better trees than wQMC, and usually achieves better quartet scores. While prior studies also reported this observation [9, 38], this study confirms this on a broad range of model conditions and weighting schemes.

Next, we observed that the estimated species trees were significantly impacted by the choice of input gene tree distribution (e.g., BestML or non-parametric bootstrapping (BS) or Bayesian trees). Quartet-based methods on BestML trees may produce highly accurate trees (very often better than those on BS trees) when estimated gene trees are reasonably accurate. However, using a distribution of trees (estimated using Bayesian MCMC techniques) instead of a point estimate BestML tree for each gene substantially improved the performance of wQFM (wQFM-GTF-MB). Similarly, the branch supports estimated based on Bayesian tree distributions make weighted ASTRAL more accurate than ASTRAL. Thus, appropriate weighting based on gene tree distributions can reduce the incongruence often observed across different species tree inference pipelines. However, wQFM-GTF-MB was consistently better than wASTRAL. So, leveraging distribution of trees for each gene (preferably with Bayesian MCMC techniques), generating weighted quartets based on them, and estimating species trees using wQFM may produce substantially better trees than ASTRAL and wASTRAL.

We also examined the performance of SVDquartets and BUCKy and the impact of using quartets weighted based on SVD score (computed by SVDquartets) or concordance factors (computed by BUCKy). We found BUCKy to be more accurate than SVDquartets and to match or outperform ASTRAL in many model conditions. On the other hand, the species trees estimated by wQFM using weighted quartets produced by SVDquarets and BUCky are often reasonably accurate but were not among the best ones identified in this study.

Taken together, the results on biological and simulated data demonstrate that the wQFM can produce the best known species tree accuracy when Bayesing gene tree distributions are given as input. Notably, in many cases, the improvement of wQFM-GTF-MB over ASTRAL (when given the same MB-estimated tree distribution as input) is substantially large (Figure 10, 11). ASTRAL has evolved through versions and has been the most accurate and widely used method over the past decade. Therefore, any improvement over ASTRAL is a notable advancement. In this context, the substantial improvement achieved by wQFM-GTF-MB is remarkable and underscores the broader impact and a clear benefit of wQFM over ASTRAL that wQFM can be used outside the context of gene tree estimation and can take weighted quartets computed from various sources.

Taking all these observations into consideration, we make the following recommendations. First, if gene trees are well estimated, we can then use BestML trees with existing summary methods (ASTRAL, wQFM, BUCKy, SVDquartets). However, under challenging model conditions with limited number of erroneous estimated gene trees, appropriate methods need to be chosen as their relative performance varies depending on the complexity of the dataset/model conditions. Under such practical model conditions, utilizing distributions of trees (preferably with Bayesian MCMC techniques) may result in higher species tree accuracy.

Second, we recommend considering multiple approaches to species tree estimation, such as wQFM-GTF, wQFM-GTF-MB, ASTRAL, and wASTRAL. When these analyses yield conflicting results, examining the underlying reasons for the disagreement can help identify which analysis is likely to be more reliable or indicate the need for additional data, such as more genes or taxa, or improved data quality (e.g., more accurate alignments and more accurate gene trees).

While this study addresses a broad range of challenging model conditions and considers a variety of techniques for quartet distribution generation and quartet-based species tree inferences, it has several limitations and potential areas for extension. This study is limited to small-and moderate-sized datasets, excluding large datasets due to the computational demands of Bayesian techniques such as MrBayes. Specifically, this study explored performance on a large number of genes but did not explore large numbers of taxa, nor under conditions with missing data (i.e., gene tree with missing taxa). Additionally, this study does not include co-estimation techniques like BEST [45], *BEAST [46] due to their high computational requirements.

This study evaluated summary methods using a single maximum likelihood (ML) tree estimate for each gene, a set of ML gene trees estimated for the bootstrap replicates of each gene, or a set of trees for each gene generated using Bayesian MCMC sampling. Future studies need to consider other approaches for leveraging distributions of trees for each gene. Specifically, investigating the performance of multi-locus bootstrapping (MLBS) [51], including resampling both sites and genes, would be an interesting research direction.

Finally, this study, like any other, did not capture all the complexities inherent in real biological data. We did not investigate scenarios where gene tree discordance may have been influenced by other biological factors, such as gene duplication and loss [1, 52, 53], gene flow [54], recombination [55], horizontal gene transfer, or hybridization [56]. Other factors, such as deep versus shallow radiations, heterotachy, mistaken homology, and violations of the model of sequence evolution, should be adequately studied in future studies.

## Supporting information

Supplementary results and data

## Data Availability

The scripts used to perform the experimental study are available at https://github.com/navidh86/ quartet-inference-comparative-study.

Datasets are available at DRYAD repository: 10.5061/dryad.wstqjq2wn. Link for peer-review:

https://datadryad.org/stash/share/fQW9FYjFMHarc7X40fyyjoDYU5cRFhweQRPKuDKRTaU.

## Additional files

**Supplementary materials SM1:** Supplementary results and data.

