## Supplementary results and data for "Leveraging weighted quartet distributions for enhanced species tree inference from genome-wide data"

These supplementary materials present additional results, supplementary figures, and tables.

### 1 Additional Results

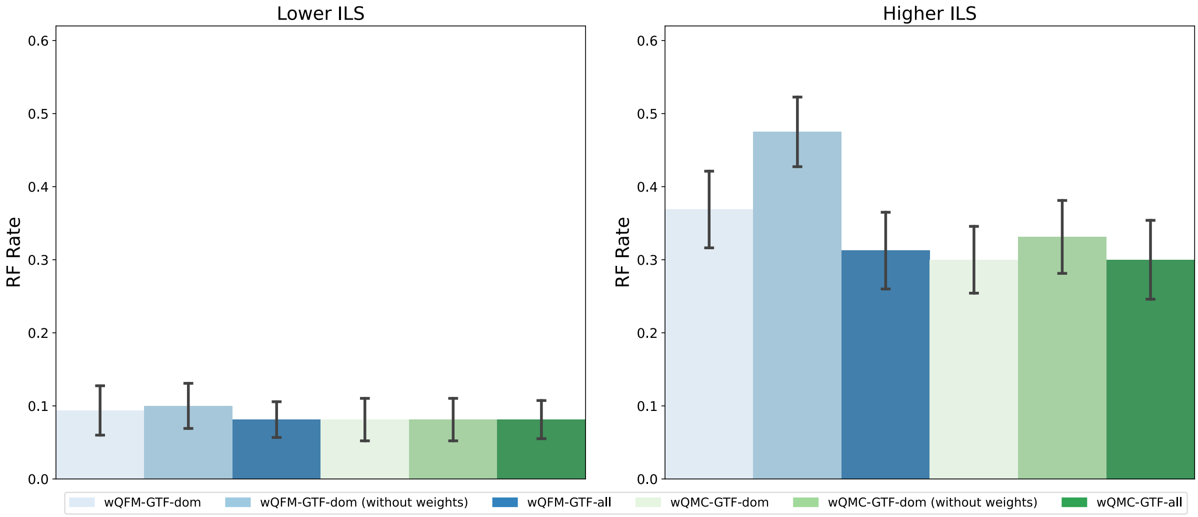


Figure S1: **RQ1 (Experiment 1): Results on the 11-taxon dataset.** Comparison of performance when using all quartets with weights and using only the dominant quartets (with and without weights).

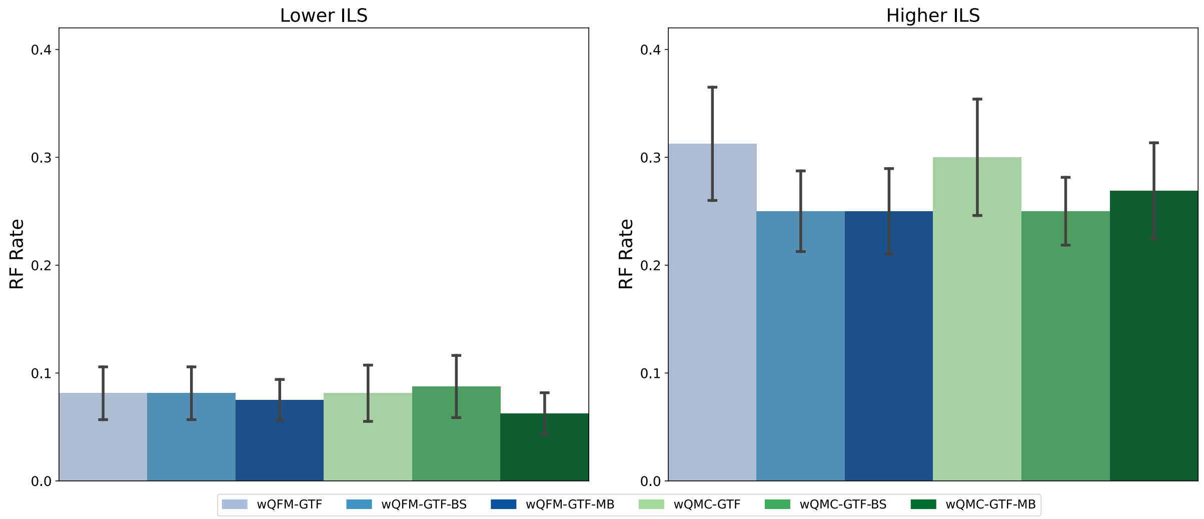


Figure S2: **RQ1 (Experiment 2): Results on the 11-taxon dataset.** We compare methods utilizing weighted quartets generated from BestML gene trees (GTF), bootstrap distribution of gene trees (GTF-BS), and Bayesian distribution of gene trees (GTF-MB).

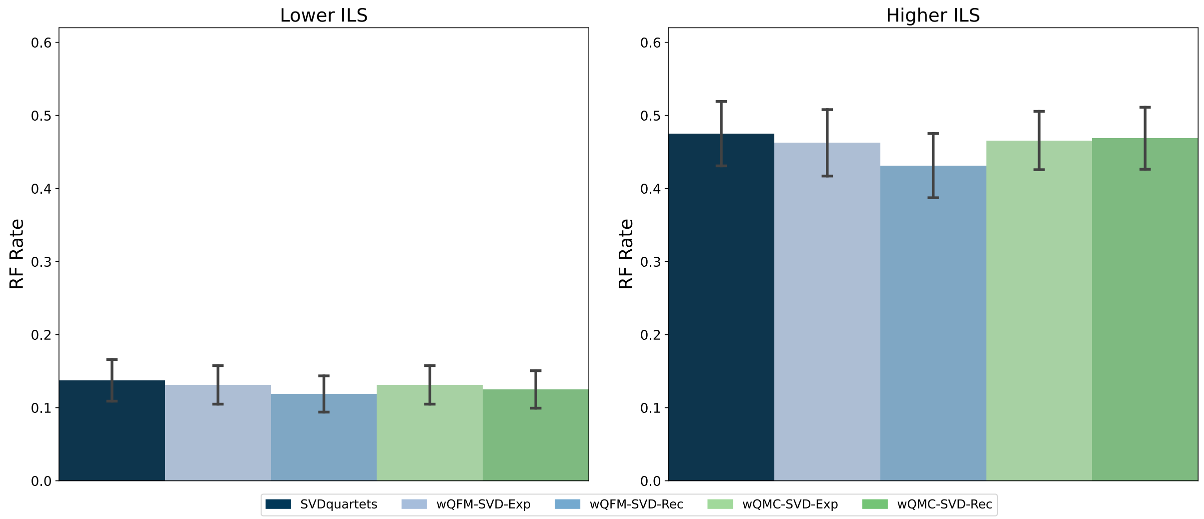


Figure S3: **RQ1 (Experiment 3): Results on the 11-taxon dataset.** Performance comparison between unweighted and weighted (both exponential and reciprocal) quartets generated by SVDquartets. The unweighted quartets are amalgamated by QFM, whereas the weighted ones are processed by both wQFM and wQMC.

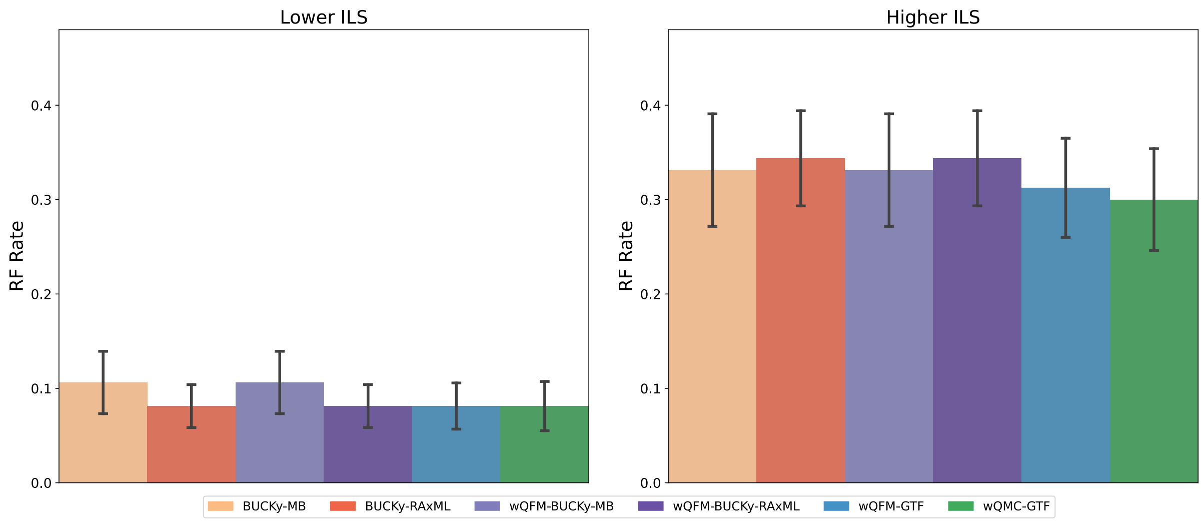


Figure S4: **RQ1 (Experiment 4): Results on the 11-taxon dataset.** Comparison of various BUCKy-based and ML-based methods. We also included the methods wQFM-GTF and wQMC-GTF.

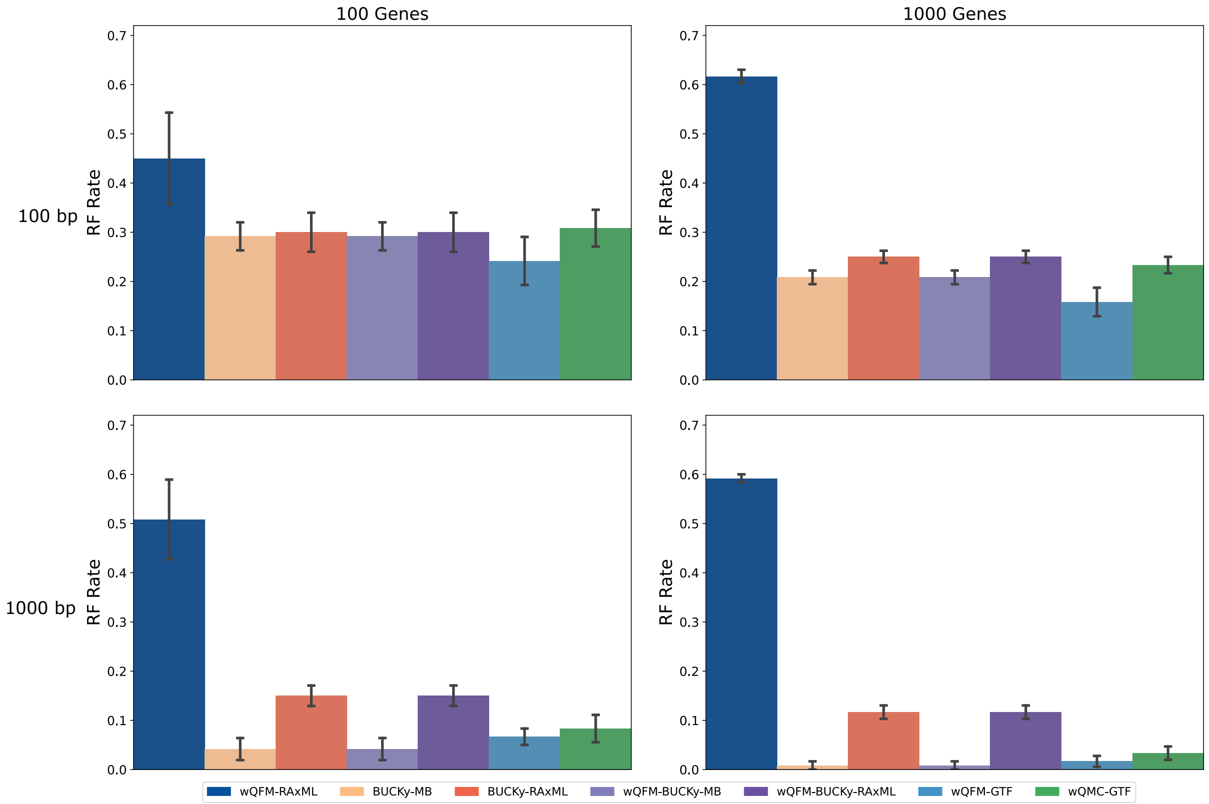


Figure S5: **RQ1 (Experiment 4): Results on the 15-taxon dataset.** Comparison of various BUCKy-based and ML-based methods with wQFM-RAxML added. We also included wQFM-GTF and wQMC-GTF.

Table S1: Quartet scores of wQFM and wQMC for estimated and true gene trees in different model conditions. The highest scores have been highlighted in bold, and those closest to the true score have been italicized.

| **Taxa** | **Model Condition** | **Estimated Gene Trees** | | | **True Gene Trees** | | |
|---|---|---|---|---|---|---|---|
| | | **wQFM-GTF** | **wQMC-GTF** | **True Tree** | **wQFM-GTF** | **wQMC-GTF** | **True Tree** |
| 11 | lower-ILS | *40412* | **40421** | 40301 | 50737 | ***50809*** | 51252 |
| | higher-ILS | *26680* | **26702** | 26372 | 29268 | ***29295*** | 29982 |
| 15 | 100gene-100bp | *69776* | **69930** | 69307 | ***82708*** | 82045 | 84634 |
| | 100gene-1000bp | *82129* | **82166** | 82099 | ***84437*** | 84285 | 84634 |
| | 1000gene-100bp | *692949* | **693656** | 690268 | ***834890*** | 827970 | 844184 |
| | 1000gene-1000bp | *817993* | **818022** | 817937 | ***843791*** | 843362 | 844184 |
| 37 | 1X-200-50 | *7523379* | **7523568** | 7517955 | ***11707475*** | 11659108 | 11744078 |
| | 1X-200-250 | *10568957* | **10569443** | 10565291 | ***11735692*** | 11732415 | 11744078 |
| | 1X-200-500 | *11271425* | **11271905** | 11267990 | ***11739571*** | 11738991 | 11744078 |
| | 1X-200-1000 | *11586354* | **11586641** | 11584969 | 11743720 | ***11743809*** | 11744078 |
| | 0.5X-200-500 | **10006593** | *10006510* | 10003929 | ***10311452*** | 10310345 | 10311622 |
| | 2X-200-500 | *11947020* | **11947266** | 11946371 | ***12565347*** | 12565281 | 12570342 |
| | 1X-100-500 | *5639605* | **5640131** | 5636883 | ***5871208*** | 5870844 | 5874698 |
| | 1X-500-500 | *28171681* | **28171698** | 28171385 | ***29363114*** | 29362939 | 29364013 |

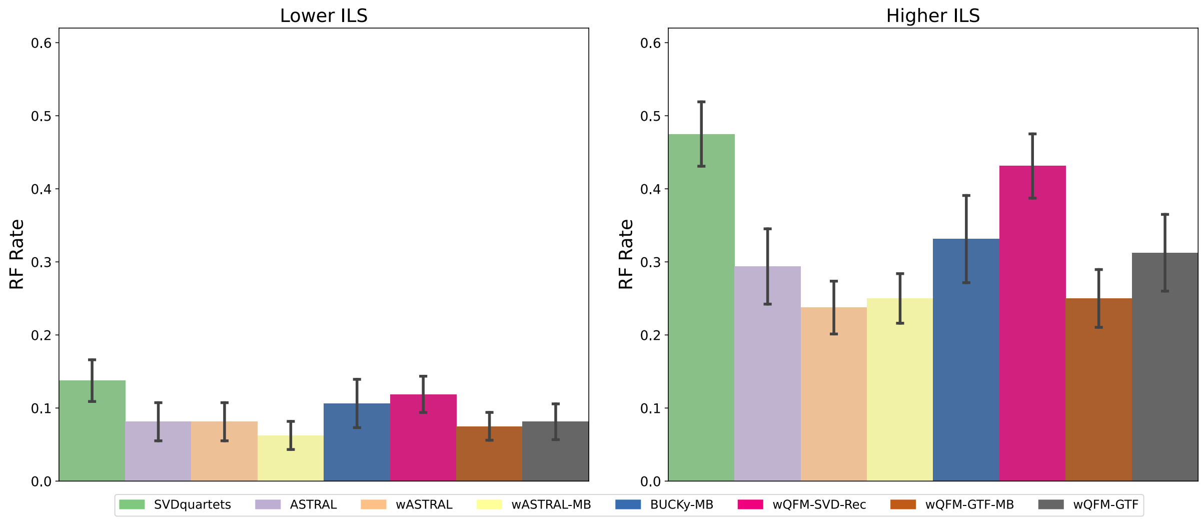


Figure S6: **RQ3: Results on the 11-taxon dataset.** We compare the best methods from previous experiments: wQFM-GTF, wQFM-GTF-MB, wQFM-SVD-Rec with ASTRAL, BUCKy, and SVDquartets. ASTRAL's weighted counterpart, configured to utilize branch supports as weights, was also analyzed. Branch supports were estimated from both non-parametric RAxML bootstrapping (wASTRAL) and Bayesian MCMC sampling (wASTRAL-MB).
